# Early locus coeruleus tauopathy features potassium channel dysfunction and is attenuated by Kv7 activation

**DOI:** 10.64898/2026.11.19.767760

**Authors:** Yi-Ci Zhang, Yi-Ge Wang, Xue-Ting Zhang, Peng-Yue Chen, Kang Tan, Zi-Yue Zhou, Xia Min, Wen Zeng, Zhuo-Hao He, Hua-Tai Xu, Kai-Wen He

**Author notes:** **Correspondence:** Kai-Wen He.

## Abstract

**Background:** The locus coeruleus (LC) accumulates abnormal tau early in Alzheimer’s disease, yet the mechanisms linking early tauopathy to neuronal dysfunction and the potential to modify subsequent pathology remain unclear. We investigated potassium-dependent excitability control, associated molecular changes and Kv7-targeted intervention.

**Methods:** We combined electrophysiology, single-cell RNA sequencing, immunohistochemistry and sleep–electroencephalographic recordings in P301S tau transgenic (PS19) mice and wild-type mice receiving local LC inoculation of synthetic human tau fibrils. Pharmacological channel modulation and glycogen synthase kinase-3β (GSK-3β) inhibition assessed pharmacological modulation of the electrophysiological phenotype. Chronic retigabine treatment assessed pathological outcomes after unilateral locus coeruleus fibril inoculation in young PS19 mice.

**Results:** Young, but not aged, PS19 mice exhibited elevated tonic firing and intrinsic excitability of locus coeruleus neurons, associated with pre-tangle phosphorylated tau accumulation and reduced low-threshold potassium current density. Exploratory single-cell profiling identified candidate expression changes in marker-defined noradrenergic neurons: increased stress-associated gene expression coincided with reduced expression of neurofilament, neuronal-maintenance and potassium-channel genes, including *Kcnq3* and *Kcnd1*. Increased nuclear RE-1 silencing transcription factor (REST) and reduced Kv7.2 and Kv7.3 immunoreactivity suggested a candidate transcriptional mechanism associated with channel dysfunction. Local fibril inoculation reproduced the physiological abnormalities in wild-type mice, with hyperexcitability depending on endogenous tau. Combined Kv4/Kv7 modulation attenuated hyperexcitability, while GSK-3β inhibition reduced tau phosphorylation, increased low-threshold potassium currents and lowered tonic firing. Both models showed increased dark-phase wakefulness and state-dependent electroencephalographic power changes. Chronic treatment with the Kv7 opener retigabine reduced bilateral LC AT8-positive tau burden and was associated with higher tyrosine hydroxylase (TH)-positive neuronal density after unilateral seeding.

**Conclusions:** Early LC tauopathy combines impaired potassium-channel-dependent excitability control with transcriptional changes consistent with cellular stress and reduced neuronal maintenance. Kv7 activation attenuates seeded tau pathology, supporting excitability regulation as a candidate intervention strategy during early tauopathy.

## INTRODUCTION

Neurofibrillary tangles (NFTs) composed of aggregated, abnormally phosphorylated tau are a defining neuropathological feature of Alzheimer’s disease (AD). However, pre-tangle tau accumulation can precede overt cognitive impairment by decades. The locus coeruleus (LC) — a pontine nucleus that provides widespread noradrenergic innervation — is among the earliest regions to accumulate abnormally phosphorylated tau. Human neuropathological studies have identified LC pre-tangle pathology in young individuals that progressively accumulates across aging and AD stages before substantial cortical tau pathology [1, 2]. As AD progresses, the LC undergoes substantial neuronal loss [3], and lower LC structural integrity is associated with greater cortical tau burden and cognitive decline [4]. These observations implicate the LC in early AD-associated processes. The functional consequences of this prolonged period of early LC tau accumulation remain incompletely understood.

The mechanisms underlying LC vulnerability in AD and other neurodegenerative diseases remain incompletely understood. Autonomous firing and activity-dependent calcium entry can generate mitochondrial oxidant stress, providing a potential physiological basis for this vulnerability [5]. Molecular studies have also identified differences in antioxidant pathways and cholesterol homeostasis between the LC and neuronal populations relatively resistant to AD- associated degeneration [6]. Nevertheless, how early tau accumulation relates to molecular changes in LC neurons, and how these changes affect their physiology, remains unclear. In particular, coordinated alterations in stress responses, neuronal maintenance and ion-channel regulation could influence LC function before overt degeneration. Linking these molecular features to neuronal excitability is therefore important for understanding the early functional consequences of LC tau pathology.

Such functional changes could have consequences for sleep and arousal. LC activity regulates sleep–wake transitions and cortical dynamics, with optogenetic manipulation demonstrating a frequency-dependent causal relationship between LC firing and transitions from sleep to wakefulness [7]. Beyond promoting wakefulness, LC-derived norepinephrine exhibits infraslow fluctuations during non-rapid eye movement (NREM) sleep that coordinate microarousals, spindle occurrence and transitions toward rapid eye movement (REM) sleep [8, 9]. Altered LC excitability could therefore affect both sleep duration and sleep microarchitecture. Although sleep disturbances are detectable in preclinical AD [10], whether early LC tau pathology contributes to these abnormalities remains unresolved.

Changes in LC activity may also influence subsequent tau pathology. In PS19 mice, inoculation of synthetic tau fibrils into the LC induces tau pathology in anatomically connected regions [11]. Experimental increases in neuronal activity enhance tau release and propagation [12], while sleep deprivation increases extracellular tau and promotes tau spreading in mice [13]. These findings raise the possibility of a reciprocal relationship in which early tau-associated LC dysfunction alters sleep regulation and neuronal activity, potentially influencing further tau accumulation. Whether early LC tau pathology alters intrinsic excitability, which molecular changes accompany this dysfunction, and whether targeting excitability modifies subsequent pathology remain open questions.

Here, using young PS19 mice and wild-type mice with local LC tau fibril inoculation, we identified early tau-associated LC hyperexcitability accompanied by reduced low-threshold potassium currents and altered sleep–wake organization. Molecular profiling nominated stress- and cell maintenance-associated transcriptional changes and reduced potassium-channel expression, including *Kcnq3*, alongside increased nuclear REST. Pharmacological modulation of potassium channels or inhibition of tau phosphorylation attenuated LC hyperexcitability, while chronic administration of retigabine reduced bilateral LC AT8 burden after unilateral fibril inoculation. These findings implicate potassium channel dysfunction in early tau-associated LC abnormalities and identify neuronal excitability as a potential early intervention target.

## RESULTS

### LC neurons exhibit elevated tonic firing in young but not older PS19 mice

LC neurons are intrinsically active pacemaker cells whose tonic firing contribute to the regulation of arousal and brain state [7, 14]. To examine how LC activity differs across stages of tauopathy, we measured spontaneous firing in acute brainstem slices from PS19 mice and age-matched wild-type (WT) littermates.

We first examined 10-month-old mice, a stage characterized by robust tau pathology and neuronal loss within the LC [15]. Histological analysis revealed AT8-positive tau pathology and increased AT8 immunoreactivity in TH-positive LC neurons, accompanied by a marked reduction in TH-immunoreactive neuron density relative to WT littermates (Fig. 1A–D). Despite these pathological changes, recorded LC neurons from PS19 mice exhibited spontaneous firing rates spanning a similar range to those of WT neurons, with no statistically significant difference in mean firing rate or resting membrane potential (Fig. 1E–G). Thus, advanced pathology and reduced TH-positive cell density were not accompanied by elevated tonic firing in the surviving LC neurons sampled (Fig. 1H).

**FIGURE 1.**
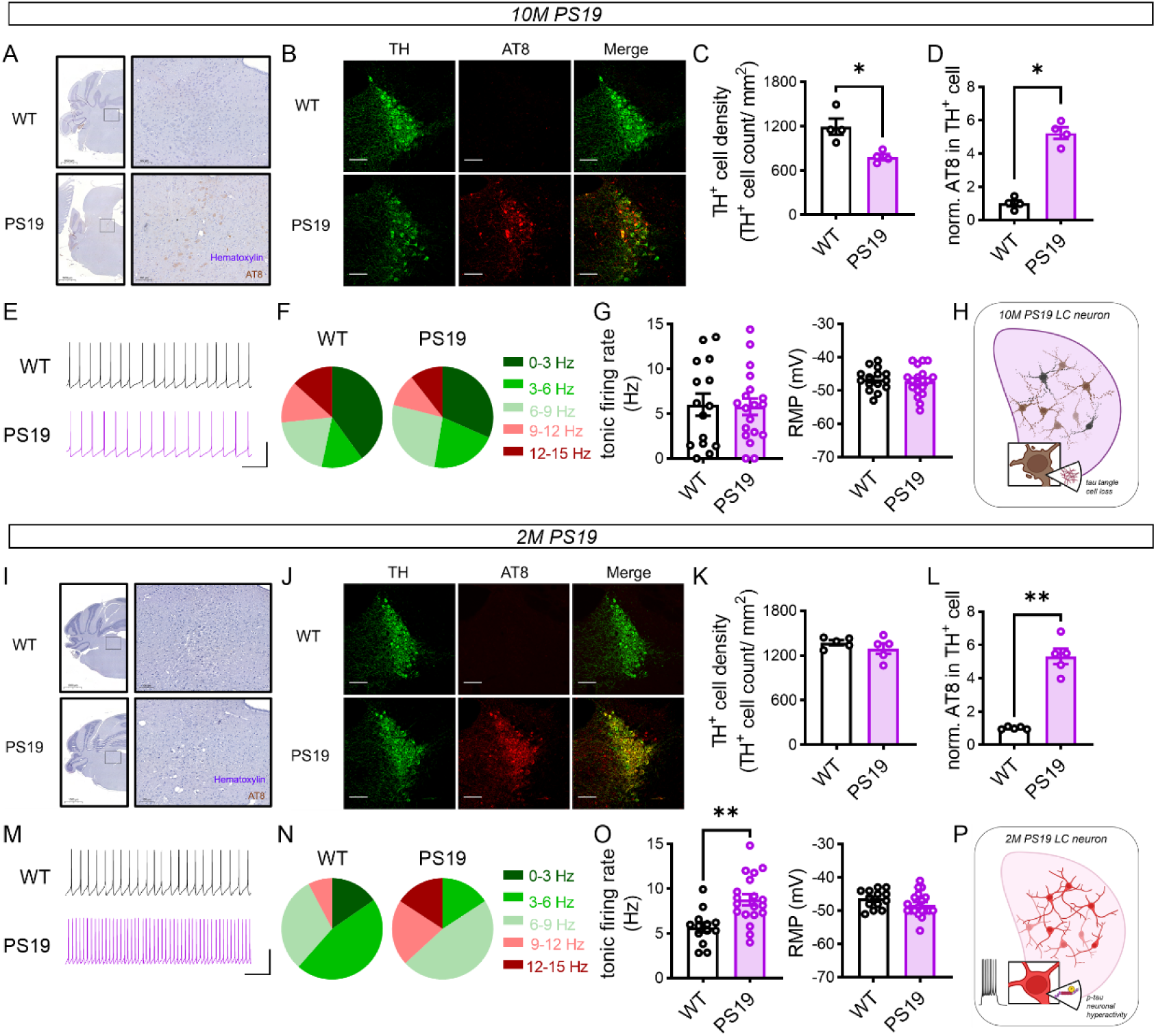
Locus Coeruleus neurons are hyperactive in 2-month-old PS19 mice. (A) Representative images and enlarged LC area stained with hematoxylin and AT8 from 10-month-old PS19 and WT mice. Boxes indicate the enlarged regions. (Scale bars, 1000 μm (overview) and 100 μm (enlargement)). (B-D) Immunostaining of the LC in 10-month-old WT and PS19 mice. (WT: n = 4 mice; PS19: n = 4 mice) (B) Representative LC images showing tyrosine hydroxylase (TH, green), AT8 (red), and merged channels. (Scale bar, 100 μm) (C) Density of TH-positive LC neurons, (p = 0.0286). (D) AT8 signal in TH-positive LC neurons, normalized to WT (p = 0.0286). (E-G) *Ex vivo* recordings of LC neurons from 10-month-old WT and PS19 mice. (WT: 15 cells from 4 mice; PS19: 19 cells from 4 mice) (E) Representative traces of recording (Scale bar, 600 ms, 25pA) (F) Distribution of firing rate. (G) Tonic firing rate (left) and resting membrane potential (RMP; right). (H) Schematic summary of 10-month-old PS19 mice LC neurons. (I) Representative images and enlarged LC area stained with hematoxylin and AT8 from 2-month-old PS19 and WT mice. Boxes indicate the enlarged regions. (Scale bars, 1000 μm (overview) and 100 μm (enlargement)). (J-L) Immunostaining of the LC in 2-month-old WT and PS19 mice. (WT: n = 5 mice; PS19: n = 5 mice) (J) Representative LC images showing tyrosine hydroxylase (TH, green), AT8 (red), and merged channels. (Scale bar, 100 μm). (K) Density of TH-positive LC neurons. (L) AT8 signal in TH-positive LC neurons, normalized to WT (p = 0.0079). (M-O) *Ex vivo* recordings of LC neurons from 2-month-old WT and PS19 mice. (WT: 13 cells from 4 mice; PS19: 19 cells from 4 mice) (M) Representative traces of recording (Scale bar, 600 ms, 25pA) (N) Distribution of firing rate. (O) Tonic firing rate (left; p = 0.0012) and resting membrane potential (RMP, right). (P) Schematic summary of 2-month-old PS19 mice LC neurons. Data are shown as mean ± SEM in Table S2. Mann–Whitney tests were performed.

We next asked whether LC activity was altered at an earlier stage, before detectable changes in neuronal density. In 2-month-old PS19 mice, AT8 immunoreactivity was increased in TH- positive LC neurons, without overt AT8-positive inclusions detectable by the histological assessment (Fig. 1I–L). TH-positive cell density did not differ significantly between genotypes (Fig. 1K). Neither glial fibrillary acidic protein (GFAP) intensity nor ionized calcium-binding adaptor molecule 1 (Iba1) intensity differed significantly in the LC region (Fig. S1C–D), consistent with previous reports of early tau accumulation preceding overt pathology [16].

In contrast to the older cohort, LC neurons from 2-month-old PS19 mice exhibited significantly higher tonic firing rates than age-matched WT neurons, without a significant change in resting membrane potential (Fig. 1M–O). The firing-frequency distribution was shifted descriptively toward higher frequencies: approximately 70% of WT neurons fired below 6 Hz, compared with fewer than 20% of PS19 neurons, while neurons firing at ≥ 9 Hz were more frequent in PS19 mice (Fig. 1N).

Together, these cross-sectional comparisons identify elevated tonic firing as an early LC phenotype in PS19 mice, accompanying increased tau phosphorylation before detectable reductions in TH-positive cell density or changes in the glial staining measures examined (Fig. 1P). The absence of elevated firing in the older cohort indicates that LC hyperactivity is not uniformly expressed across disease stages and motivates investigation of the mechanisms underlying early excitability dysregulation.

### Young PS19 LC neurons are hyperexcited mediated by impaired low-threshold potassium current density

Neuronal hyperactivity can result from changes in the excitatory and/or inhibitory synaptic inputs [17] or altered excitability [18]. To investigate the mechanism underlying increased tonic firing in young PS19 mice, we first examined excitatory and inhibitory synaptic transmission onto LC neurons. Neither the frequency nor amplitude of spontaneous excitatory postsynaptic currents (sEPSCs) differed significantly between genotypes (Fig. 2A). By contrast, miniature inhibitory postsynaptic current (mIPSC) frequency was increased in PS19 neurons, whereas amplitude was unchanged (Fig. 2B). Thus, the measured synaptic responses did not reveal changes that could readily account for the elevated tonic firing.

**FIGURE 2.**
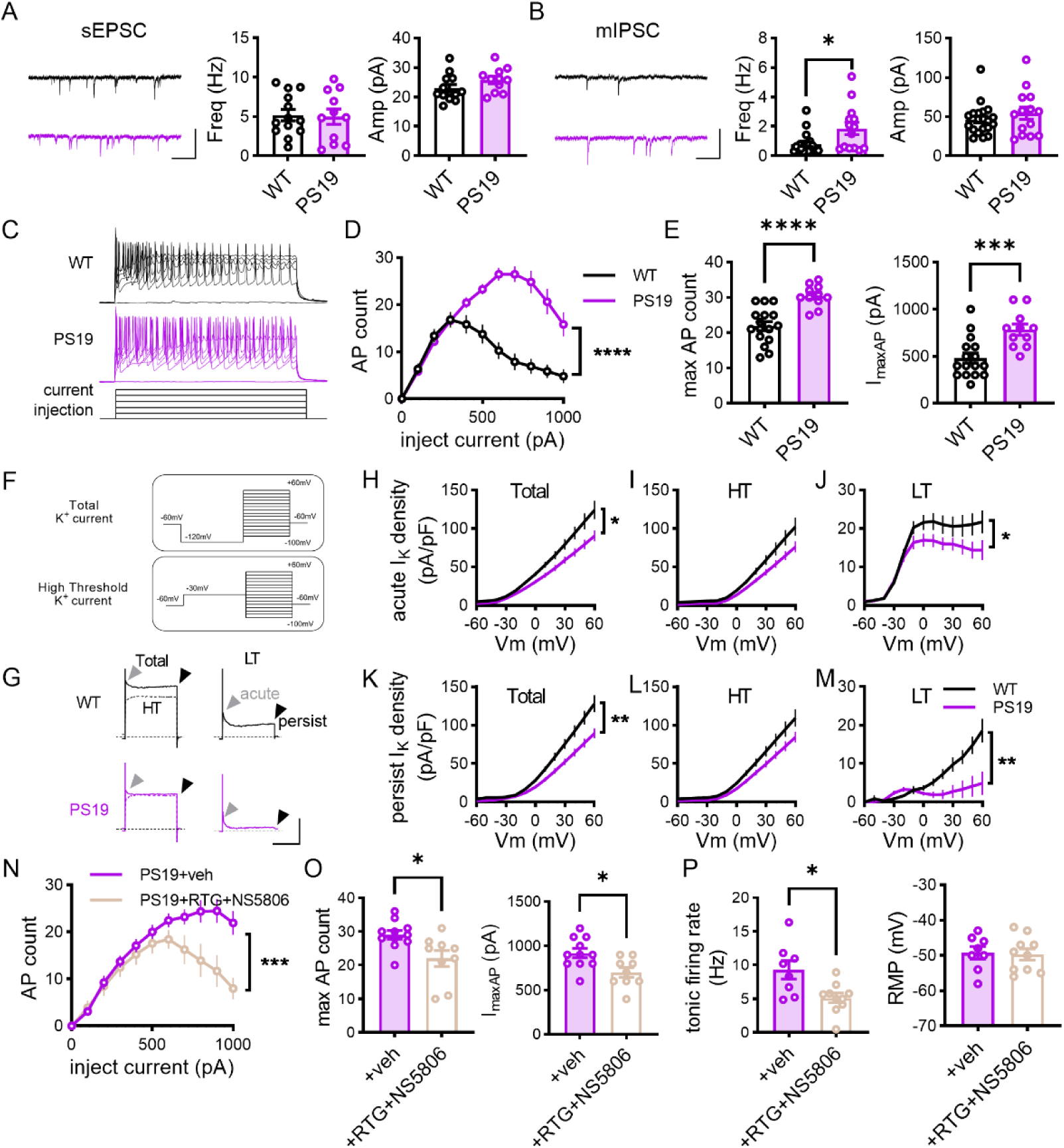
Intrinsic hyperexcitability and reduced voltage-gated potassium conductance in LC neurons of young PS19 mice. (A) Comparison of sEPSC (WT: 13 cells from 3 mice; PS19: 11 cells from 3 mice). Left panel, example sEPSC traces. Middle and right panels, quantification of the sEPSC frequency (Middle) and amplitude (Right) (Scale bars: 50 pA, 250 ms) (B) Comparison of mIPSC (WT: 20 cells from 4 mice; PS19: 14 cells from 4 mice). Left panel, example mIPSC traces. Middle and right panels, quantification of the mIPSC frequency (Middle, p = 0.0167) and amplitude (Right) (Scale bars: 50 pA, 250 ms) (C-E) Comparison of neuronal excitability by injecting current steps (WT: 16 cells from 4 mice; PS19: 11 cells from 4 mice). (C) Representative traces evoked by injecting 500 ms current steps. (D) Current-spiking relationship of LC neurons. (F (1, 25) = 38.77, p < 0.0001) (E) Quantification of maximum AP count (Left panel, p < 0.0001) and currents required to generate the maximal firing (I_maxAP_, Right panel, p = 0.0009). (F) Diagram depicting protocols measuring the total voltage-gated potassium current (I_k,_ upper panel) and high threshold (HT) I_k_ (lower panel). (G-M) Comparison of Kv channel conductance of LC neurons (WT: 13 cells from 3 mice; PS19: 10 cells from 3 mice). (G) Right panels, example traces of total (solid traces) and HT (dotted traces) I_k_. Left panels, example traces of low threshold (LT) I_k_. Grey arrowheads: acute I_k_. Black arrowheads: persistent I_k_. (Scale bar: 2000 pA, 300 ms). (H-M) V-I_k_ curves of acute (H-J) and persistent (K-M) I_k_ density. Left panels, total I_k_; Middle panels, HT I_k_; Right panels, LT I_k_. (H: p = 0.0245; J: p = 0.0464. K: p = 0.0079; M: p = 0.0068) (N-P) Effect of co-application of retigabine (RTG) and NS5806 on neuronal excitability (N-O, Veh: 11 cells from 3 mice; RTG+NS5806: 9 cells from 3 mice) and tonic firing rate (P, Veh: 8 cells from 3 mice; RTG+NS5806: 10 cells from 3 mice) of LC neurons of PS19 mice. (N) Current-spiking relationship. (F (1, 18) = 15.46, p = 0.0010) (O) Maximum AP count (left; p = 0.0103) and I_maxAP_ (right; p = 0.0153). (P) Spontaneous tonic firing rate (left; p = 0.0266) and RMP (right) Data are shown as mean ± SEM in Table S3. Mann–Whitney tests were performed. Two-way ANOVA with Sidak post hoc multiple-comparisons test were used to compare I-V curves and I-O curves.

We next examined neuronal excitability using depolarizing current injections. LC neurons from young, but not older (Fig. S2A, B) PS19 mice generated more action potentials at stronger current steps, required a much larger injected current (I_maxAP_) to reach a higher maximal spike count (Fig. 2C-E). These findings were consistent with reduced susceptibility to depolarization block. Action potential threshold, rheobase and input resistance did not differ significantly between genotypes (Fig. S2C-E). Analysis of spontaneous tonic AP properties revealed a trend toward reduced afterhyperpolarization (AHP) amplitude (Fig. S2F-G), and a smaller AHP area and shorter recovery times (Fig. S2H–J). Together, these changes indicated LC neurons in young PS19 mice were hyperexcitable with altered post-spike voltage recovery.

To determine whether reduced outward potassium currents contributed to this phenotype, we measured voltage-gated potassium currents (I_k_) using two conditioning potentials (Fig. 2F). Currents recorded after conditioning at −120 mV were designated total currents, whereas those recorded after conditioning at −30 mV were designated high-threshold (HT) currents [19]. The low-threshold (LT) component was estimated by subtracting the latter from the former (Fig. 2G; see Methods). Both acute (fast-kinetic, Fig. 2G, gray arrow head) and persist (slow-kinetic, Fig. 2G, black arrow head) I_k_ amplitudes were quantified. LC neurons from PS19 mice showed a significant reduction in total I_k_, including both acute and persistent components, with reductions detected in the LT component but not the HT component (Fig. 2H-M).

These deficits prompted us to examine Kv4 and Kv7 channels as candidate contributors. Kv4 channels can mediate rapidly inactivating A-type currents, whereas Kv7 channels mediate slowly activating, non-inactivating M-type currents[19]. Co-application of Kv4 channel modulator NS5806 and the Kv7 channel opener retigabine effectively mitigated the hyperexcited phenotypes, including decreased maximal firing rate and reduced current required for generating the maximal firing (Fig. 2N-O). Combined treatment also lowered tonic firing without a significant change in resting membrane potential (Fig. 2P). Complementing these findings, we previously showed in wild-type LC neurons that separate inhibition of Kv7 and Kv4 increased neuronal excitability and firing [20].

Together, these findings support a contribution of reduced LT K^+^ currents to early LC hyperexcitability in PS19 mice and identify Kv4 and Kv7 channels as candidate components of the impaired excitability-control mechanism.

### Local tau fibril inoculation induces tau hyperphosphorylation and reproduces electrophysiological abnormalities in wild-type LC neurons

Tau hyperphosphorylation and increased neuronal excitability co-occurred in LC neurons of young PS19 mice. To test whether locally induced tau abnormalities reproduce this physiological phenotype without transgenic tau overexpression, we used heparin-induced human tau preformed fibrils containing the frontotemporal lobar degeneration-associated P301L mutation (Hep-T40PL PFF, hereafter referred to as fibrils, Fig. 3A). Such fibrils have previously been shown to induce pathological tau aggregation in tau transgenic mice [11, 21–23]. Compared with the corresponding T40PL monomer (hereafter referred to as monomer), fibril exposure increased tau phosphorylation in cultured wild-type hippocampal neurons (Fig. 3B-D), supporting their use to induce tau phosphorylation in non-transgenic neurons.

**Figure 3.**
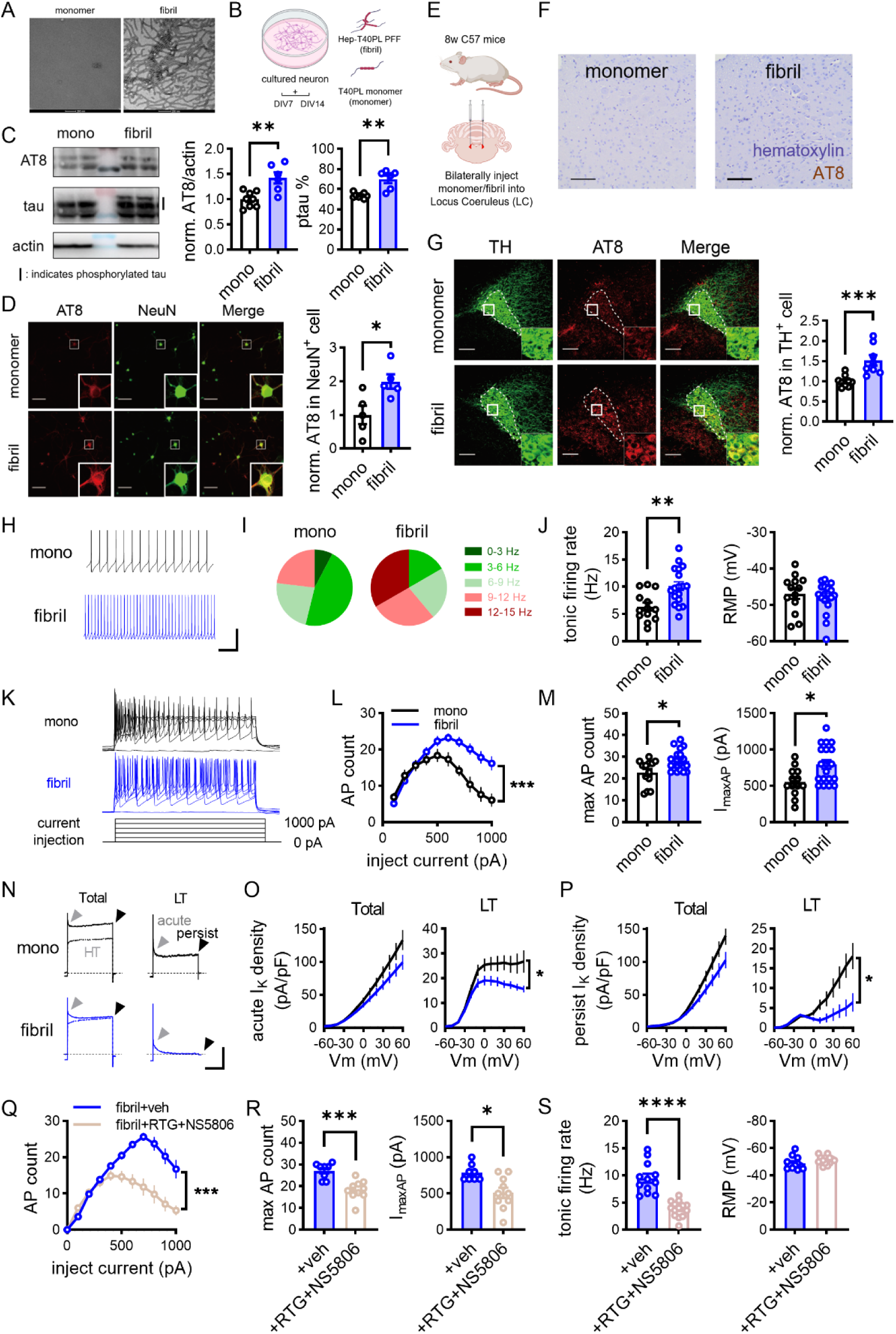
Synthetic human tau preformed fibril induces local tau hyperphosphorylation and PS19-like hyperexcitability in LC neurons of wild-type mice. (A) Representative negative staining electro-microscopy images of tau monomer (left, no heparin) and heparin-induced tau preformed fibrils (right). (Scale bar, 200 nm). (B) Schematic diagram of primary cultured neurons treated with either 1 μg Hep-T40PL-PFF (fibril) or T40PL monomer (monomer) at DIV7 for 1 week. (C) Western blot analysis of tau phosphorylation after fibril or monomer treatment of cultured neurons (mono: n = 8 wells; fibril: n = 6 wells). Left panel, representative blot images. Middle panel, quantification of AT8 (normalized to actin). Right panel, quantification of the percentage of phosphorylated tau (upper bands normalized to all bands detected by tau antibody). (AT8/actin: p = 0.0047; p-tau (%): p = 0.0027) (D) Tau phosphorylation was assessed by AT8 immunostaining in cultured neurons. Left, representative images of AT8 (red), NeuN (green), and merged channels (Scale bar, 100 µm). Right, quantification of normalized AT8 intensity in NeuN-positive cells, averaged per well (mono: n = 5 wells; fibril: n = 5 wells) (p = 0.0317) (E) Schematic diagram showing the *in vivo* fibril inoculation into wild-type mice bilateral LC. (F) Immunochemical staining of AT8 (brown) and hematoxylin (purple) revealed no trace of pathological tau in the LC regions of either monomer- or fibril-inoculated wild-type mice. (Scale bar, 100 µm) (G) Immunostaining analysis of tau phosphorylation induced by fibril inoculation. Left: Representative images showing AT8 (red) co-stained with TH (green) in LC regions (dotted line) of monomer- or fibril-inoculated wild-type mice (Scale bar, 100 µm). Right: Quantification of the normalized AT8 signal intensity per TH+ soma. (mono: n = 9 mice; fibril: n = 8 mice) (p = 0.0006) (H-J) *Ex vivo* recordings of tonic firing of LC neurons of fibril- (blue) or monomer-inoculated (black) wild-type mice. (mono:13 cells from 3 mice; fibril: 18 cells from 4 mice) (H) Representative traces of recording (Scale bar, 600ms, 25pA). (I) Distribution of firing rate. (J) Quantification of tonic firing rate (Left, p = 0.0049) and resting membrane potential (RMP, right). (K-M) LC neurons showed hyperexcitability in wild-type mice inoculated with tau fibril (mono: 13 cells from 3 mice; fibril: 19 cells from 4 mice). (K) Representative traces evoked by injecting 500 ms current steps. (L) Current-spiking relationship of LC neurons. F (1, 30) = 17.87, p = 0.0002. (M) Quantification of maximal firing rate (left, p = 0.047) and currents required to generate the maximal firing (I_maxAP_, right, p = 0.0117). (N) Example traces of total (left line), high threshold (HT, left dotted line) and low threshold (LT, right line) voltage-gated I_k_ from mono and fibril-inoculated mice LC neurons. Grey arrowheads indicate acute current. Black arrowheads indicate persist current. (Scale bar: 2000 pA, 300 ms). (O-P) V-I curves of acute (O) and persist (P) I_k_ density of total (left panels) and LT (right panels) I_k_. (mono: 12 cells from 3 mice; fibril: 13 cells from 3 mice) (O, right: p = 0.0382; P, right: p = 0.0154) (Q-S) Effect of co-application of retigabine (RTG) and NS5806 on neuronal excitability (Q-R, Veh: 8 cells from 3 mice; RTG+NS5806: 10 cells from 3 mice) and tonic firing rate (S, Veh: 12 cells from 3 mice; RTG+NS5806: 14 cells from 3 mice) of LC neurons of fibril-inoculated mice. (Q) Current-spiking relationship. (F (1, 16) = 23.75, p = 0.0002) (R) Maximum AP count (left; p = 0.0004) and I_maxAP_ (right; p = 0.0104). (S) Spontaneous tonic firing rate (left; p < 0.0001) and RMP (right) Data are shown as mean ± SEM in Table S4. Mann–Whitney tests were performed. Two-way ANOVA with Sidak post hoc multiple-comparisons test were used to compare I-V curves and I-O curves.

We next bilaterally inoculated fibrils or monomer into the LC of 8-week-old wild-type mice (Fig. 3E). One month after injection, fibrils increased AT8 immunoreactivity in TH+ LC neurons (Fig. 3G), without overt AT8-positive inclusions detectable by the histological assessment (Fig. 3F). Monomer inoculation did not increase AT8 immunoreactivity relative to phosphate-buffered saline controls (Fig. S3A). TH+ cell density and TH signal intensity did not differ between fibril- and monomer-inoculated mice (Fig. S3B). Moreover, no statistically significant increases in neuronal AT8 immunoreactivity were detected in the other brain regions examined at this time point (Fig. S3C–H). These results establish a model of predominantly local tau hyperphosphorylation without detectable changes in TH-based measures of LC neuronal integrity.

To determine whether these local tau abnormalities reproduce the physiological defects observed in young PS19 mice, we recorded from wild-type LC neurons after fibril or monomer inoculation. Fibril-inoculated neurons exhibited a higher tonic firing rate without a detectable change in resting membrane potential (Fig. 3H-J), resembling the hyperactivity observed in young PS19 mice. Intrinsically, fibril-inoculated neurons were hyperexcitable with increased maximal firing rate and reduced depolarization block (Fig. 3K-M), while other membrane properties measured did not differ significantly (Fig. S4A-B).

This increase in excitability was accompanied by reduced acute and persistent LT Kv conductance (Fig. 3N-P), paralleling the deficits in young PS19 mice. To test whether potassium channel modulation could counteract the firing phenotype, we co-applied the Kv4 channel modulator NS5806 and the Kv7 channel opener retigabine (RTG). Combined treatment mitigated the hyperexcitability (Fig. 3Q-R), lowered tonic firing rate and slightly hyperpolarized resting membrane potential (Fig. 3S), supporting a contribution of impaired potassium channel-mediated excitability control.

Finally, neither the fibril-associated increase in AT8 immunoreactivity (Fig. S4E) nor the increase in neuronal excitability was detected in tau-knockout mice (Fig. S4F), indicating that these responses require endogenous tau. Together, these findings show that local tau fibril inoculation elevates tau phosphorylation and reproduces key electrophysiological features of early PS19 tauopathy in wild-type LC neurons, which support the contribution of reduced low- threshold potassium currents to the associated hyperexcitability.

### GSK-3β inhibition reduces tau phosphorylation and attenuates LC hyperexcitability in young PS19 mice

Having established that local tau fibril inoculation reproduces key electrophysiological abnormalities of young PS19 mice, we next tested whether pharmacological reduction of tau phosphorylation was accompanied by reversal of the LC firing phenotype in PS19 mice. We employed AR-A014418, an inhibitor of glycogen synthase kinase-3β (GSK-3β) previously shown to reduce tau phosphorylation [24]. *Ex vivo* incubation of brain slices from young PS19 mice with 5 μM AR-A014418 reduced AT8 immunoreactivity (Fig. 4A). We then examined whether this biochemical response was accompanied by improved Kv-mediated control of neuronal excitability. Compared with vehicle treatment, AR-A014418 significantly increased both the acute and persistent LT voltage-gated potassium current density in LC neurons (Fig. 4B-C). Consistently, it also attenuated neuronal hyperexcitability in AR-A014418-treated PS19 LC neurons (Fig. 4D-F). AR-A014418 also reduced tonic firing, without a statistically significant change in resting membrane potential (Fig. 4G–I).

**Figure 4.**
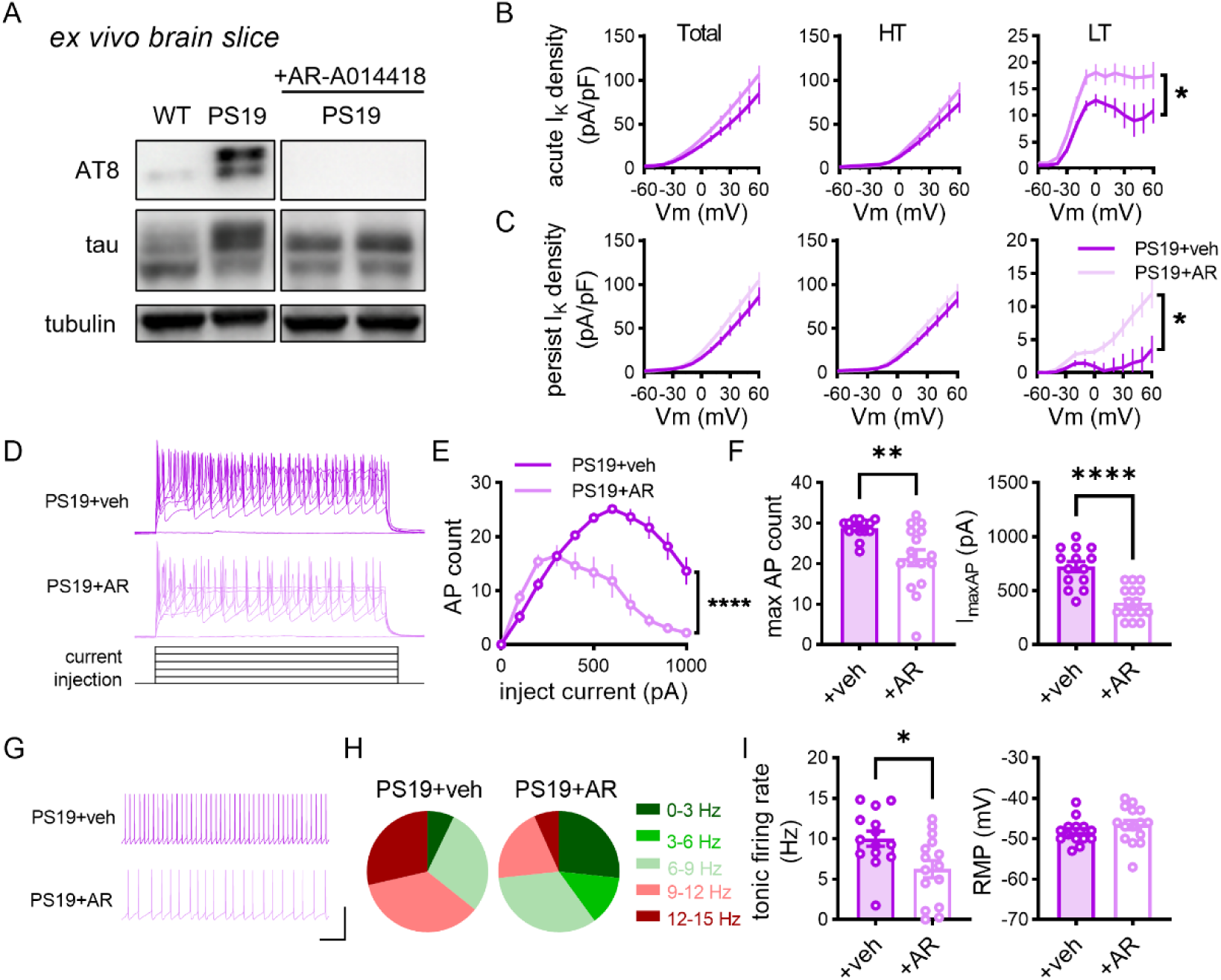
GSK-3β inhibitor restores Kv function and alleviates hyperactivity. (A) Representative AT8, tau, and tubulin immunoblots from acute WT or PS19 brain slices exposed to vehicle or 5μM AR-A014418 for 30 min. (B-C) Comparison of acute (B) and persistent (C) components of total (left panels), HT (high threshold, middle panels), and LT (low threshold, right panels) I_k_ between vehicle treated and AR-A014418 treated PS19 brain slices (Veh: 10 cells from 4 mice; AR: 16 cells from 4 mice). (B, right: p = 0.0352; C, right: p = 0.0294) (D-F) Comparison of neuronal excitability by injecting current steps (Veh: 14 cells from 4 mice; AR: 16 cells from 5 mice). (D) Representative traces evoked by injecting 500 ms current steps. (E) Current-spiking relationship of LC neurons. F (1, 28) = 27.13, p < 0.0001. (F) Quantification of maximal firing rate (left, p = 0.0027) and currents required to generate the maximal firing (I_maxAP_, right, p < 0.0001). (G-I) *Ex vivo* recordings of tonic firing of LC neurons in vehicle treated and AR-A014418 treated 2-month-old PS19 mice. (Veh: 14 cells from 4 mice; AR: 15 cells from 4 mice) (G) Representative traces of recording (Scale bar, 600ms, 25pA). (H) Distribution of firing rate. (I) Quantification of tonic firing rate (Left, p = 0.0137) and resting membrane potential (RMP, right). Data are shown as mean ± SEM in Table S5. Mann–Whitney tests were performed. Two-way ANOVA with Sidak post hoc multiple-comparisons test were used to compare I-V curves and I-O curves.

Together, these findings demonstrate that the electrophysiological abnormalities of young PS19 LC neurons are attenuated by AR-A014418. The accompanying reduction in hyperphosphorylated tau supports a relationship between tau phosphorylation and impaired excitability control.

### Single-cell profiling nominates stress-associated transcriptional changes and reduced potassium-channel expression in PS19 LC neurons

Following the identification of early LC hyperexcitability, we sought to define the transcriptional changes accompanying this phenotype in young PS19 mice. Single-cell RNA sequencing of the LC-containing tissue pooled from 8 mice each per genotype resolved 13 neuronal clusters. Cluster 13 was enriched for a panel of noradrenergic marker genes and was therefore annotated as noradrenergic (NE) neurons (Fig. 5A, B; Fig. S5). These neurons exhibited a higher immediate-early-gene enrichment score (defined as neural activity score) in PS19 than in wild-type mice (Fig. 5C, D), consistent with an activity-associated transcriptional signature accompanying the increased tonic firing observed electrophysiologically.

**FIGURE 5.**
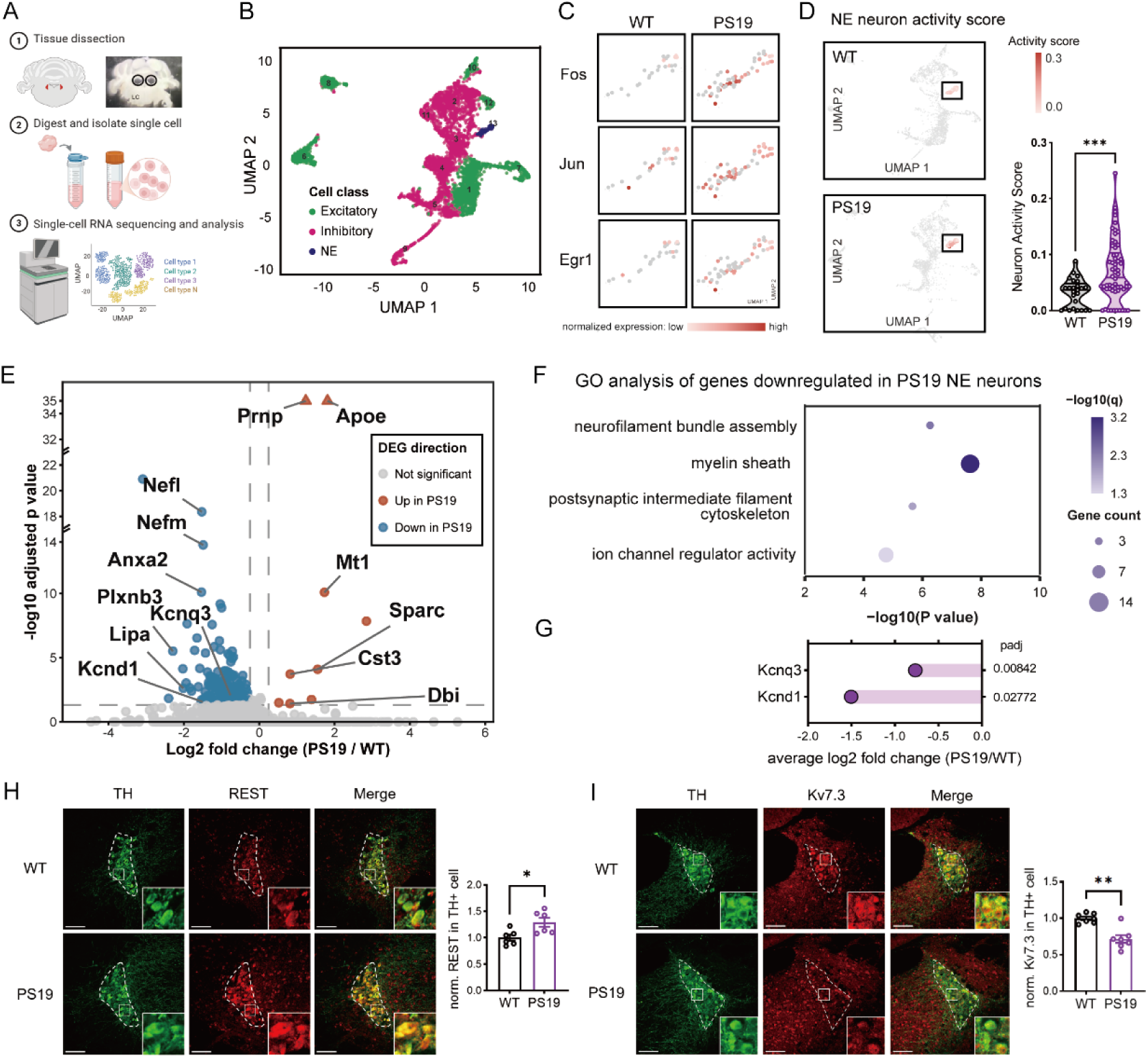
Single-cell profiling reveals activity- and Kv-related alterations in LC noradrenergic neurons in 2-month-old PS19 mice. (A) Schematic of LC dissection and single-cell RNA-sequencing workflow (WT: n = 8 mice; PS19: n = 8 mice, eight mice per genotype were pooled into one sample per genotype). (B) UMAP of neurons identified in the LC single-cell dataset, colored by neuronal class (excitatory, inhibitory, or NE). Each dot represents one cell. The NE population was assigned using the strict definition described in Methods. (C) UMAP feature plots showing expression of the activity-regulated genes *Fos, Jun* and *Egr1* in WT and PS19 neurons. (D) Left: UCell-based neuronal activity scores mapped onto the neuronal UMAP, with cluster 13 highlighted. The signature was calculated from 16 IEG/activity-regulated genes (see Methods). Right: Violin plot comparing neuron activity scores in cluster 13 neurons from WT (n = 25 cells) and PS19 (n = 58 cells). (p = 0.0008). (E) Volcano plot of differentially expressed genes in NE-like neurons from PS19 and WT mice. NE- like neurons were defined with detectable expression of at least one of *Th, Dbh, Slc6a2,* or *Slc18a2*. Genes meeting the thresholds of adjusted P ≤ 0.05 and |log₂ fold change| ≥ 0.25 are shown in red (upregulated in PS19; 9 genes) or blue (downregulated in PS19; 191 genes); other genes are gray. Selected genes are labeled. Triangles indicate genes with adjusted P ≤ 1 × 10⁻³⁵. The y-axis is displayed in three segments (0–14, 18–22, and 32–35); (F) Functional enrichment analysis of genes downregulated in PS19. The four displayed terms met the q < 0.05 threshold. The x-axis indicates −log₁₀ (P value), point size indicates gene count, and color indicates −log₁₀ (q value). (G) Differential expression of the potassium-channel genes *Kcnq3* and *Kcnd1* in NE-marker-positive neurons, shown as average log₂ fold change (PS19/WT); adjusted P values are indicated. (H) Left: Representative immunofluorescence images of TH (green) and REST (red) in the LC of WT and PS19 mice (WT: n = 6 mice, PS19: n = 6 mice). Right: Quantification of somatic REST signal in TH+ cell. (Scale bars, 100 μm) (p = 0.0152) (I) Left: Representative immunofluorescence images of TH (green) and Kv7.3 (red) in the LC of WT and PS19 mice (WT: n = 7 mice, PS19: n = 7 mice). Right: Quantification of Kv7.3 signal in TH+ cell. (Scale bars, 100 μm) (p = 0.0070) Data are shown as mean ± SEM in Table S5. Mann–Whitney tests were performed.

To broaden the population available for exploratory expression analysis, we additionally identified 579 NE-like neurons expressing at least one of the four classical NE markers (see Methods). This marker-defined population included more than 97% of cluster 13 neurons (Fig. S6A) and also exhibited an increased activity score in PS19 mice. Expression analysis identified 191 genes with lower expression and 9 genes with higher expression in PS19 NE- like neurons (Fig. 5E). Upregulated genes included *Apoe*, *Prnp* and *Mt1*, together with *Sparc* and *Cst3* (Fig. S6C), indicating changes compatible with stress responses and altered lipid handling, protease regulation and extracellular interactions. Conversely, downregulated genes were enriched for Gene Ontology terms related to neurofilament bundle assembly, myelin sheath, postsynaptic intermediate filament cytoskeleton and ion channel regulator activity (Fig. 5F). Thus, increased expression of stress-associated genes coincided with reduced expression of genes associated with neuronal structure and excitability regulation, suggesting a potentially vulnerable cellular state during early tauopathy.

Within this broader transcriptional signature, *Kcnq3* and *Kcnd1*, encoding the Kv7.3 and Kv4.1 potassium-channel subunits, respectively, were significantly downregulated (Fig. 5G). These changes provided a molecular correlate of the Kv7- and Kv4-associated current deficits. Reduced *Kcnq3* expression prompted us to examine RE1-silencing transcription factor (REST), a transcriptional repressor previously shown to regulate *Kcnq2* and *Kcnq3* expression [25], whose nuclear abundance has also been reported to increase during early tau pathology [26]. In young PS19 mice, total and nuclear REST immunoreactivity were higher in TH-positive LC neurons (Fig. 5H; Fig. S6E, F), whereas neither measure differed significantly in the NeuN- positive non-LC neurons examined (Fig. S6G). Kv7.3 immunoreactivity was also reduced in TH-positive LC neurons (Fig. 5I), consistent with the decrease in *Kcnq3* transcripts.

Together, these findings connect early LC hyperexcitability with a transcriptional signature involving stress-associated remodeling and reduced expression of structural and potassium- channel genes. The accompanying increase in nuclear REST identifies a candidate transcriptional mechanism for reduced Kv7 expression associated with early tauopathy in LC.

### Sleep and EEG alterations in fibril-inoculated wild-type mice and young PS19 mice

LC neuronal firing varies across vigilance states and contributes to the regulation of arousal and sleep–wake transitions [14, 27]. We therefore examined whether the LC hyperexcitability identified in the two tauopathy models was accompanied by altered sleep–wake organization. Electroencephalographic (EEG) and electromyographic (EMG) recordings were obtained from wild-type mice one month after bilateral LC inoculation of tau fibrils or T40PL monomer (Fig. S7A). Compared with monomer controls, fibril-inoculated mice spent more time awake and less time in non-rapid eye movement (NREM) sleep over 24 h, with these differences concentrated in the dark phase (Fig. 6A–F). During this phase, wake bouts were fewer and longer, indicating more consolidated wakefulness; rapid eye movement (REM) sleep bout counts were also reduced (Fig. S7F–I). Monomer inoculation did not alter vigilance-state proportions relative to phosphate-buffered saline controls (Fig. S7B). LC tau phosphorylation assessed after recording was positively correlated with wake percentage, whereas its negative association with NREM sleep percentage did not reach statistical significance (Fig. S7C).

**FIGURE 6.**
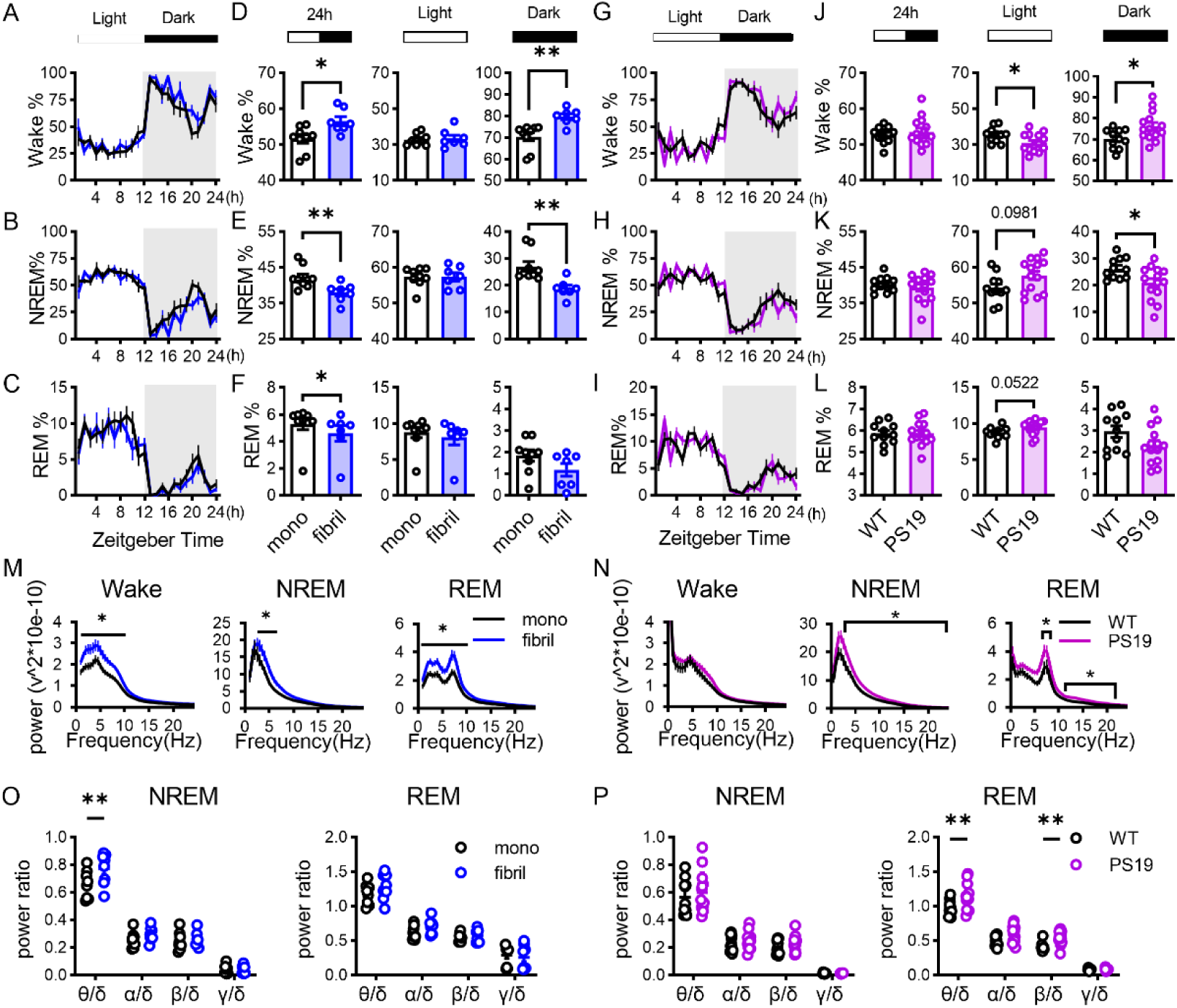
Sleep dysregulation and EEG power changes in fibril-inoculated wild-type mice and 2-month-old PS19 mice. (A–F) Percentages of time spent in wake (A), NREM (B), and REM (C) across the light/dark cycle between the fibril- and monomer-inoculated wild-type mice. Comparisons of the percentage of time spent in wake (D), NREM (E), and REM (F) during the 24-hour day (left panels), light cycle (middle panels) and the dark cycle (right panels) between the fibril- and monomer-inoculated wild-type mice. (mono: n = 9 mice; fibril: n = 8 mice). (G-L) Percentages of time spent in wake (G), NREM (H), and REM (I) across the light/dark cycle in 2-month-old PS19 mice and their WT littermates. Comparisons of the percentage of time spent in wake (J), NREM (K), and REM (L) during the 24-hour day (left panels), light cycle (middle panels) and the dark cycle (right panels) between 2-month-old PS19 mice and WT littermates. (WT: n = 11 mice; PS19: n = 14 mice) (M-N) EEG power spectrum analysis of wake (left), NREM (middle) and REM (right) in fibril- inoculated or monomer- inoculated wild-type mice (M) and 2-month-old PS19/WT mice (N). (mono: n = 9 mice; fibril: n = 8 mice; WT: n = 11 mice; PS19: n = 14 mice) (O-P) Ratios of θ-, α-, β-, and γ-band power to δ power during NREM (left panels) and REM (right panels) for the monomer/fibril (O) and WT/PS19 (P) comparisons. (mono: n = 9 mice; fibril: n = 8 mice; WT: n = 11 mice; PS19: n = 14 mice) Data are shown as mean ± SEM in Table S6. Mann–Whitney tests were performed. Multiple t-tests were used to analysis absolute power comparison. Two-way ANOVA with Sidak post hoc multiple- comparisons test were used to compare power ratios.

Young PS19 mice similarly exhibited increased wakefulness and reduced NREM sleep during the dark phase (Fig. 6G–L), together with shorter NREM bouts and fewer REM bouts (Fig. S7L, O). Their light-phase phenotype differed: wakefulness was reduced, and wake bouts were more frequent and shorter, indicating fragmentation of wakefulness during the rest phase (Fig. 6J; Fig. S7K, N). Thus, the two models shared a dark-phase shift toward wakefulness, whereas PS19 mice additionally exhibited altered light-phase sleep-wake organization.

We next examined whether these changes in vigilance-state organization were accompanied by altered EEG activity within individual states. Fibril-inoculated wild-type mice showed increased absolute spectral power during wakefulness, NREM sleep and REM sleep, whereas significant spectral differences in PS19 mice were detected during NREM and REM sleep (Fig. 6M–N). Band-to-delta power ratios further revealed an increased theta/delta ratio during NREM sleep in fibril-inoculated mice and increased theta/delta and beta/delta ratios during REM sleep in PS19 mice (Fig. 6O–P). These results identify state-dependent changes in the balance between slow and faster oscillatory activity.

Together, these findings associate local LC tau fibril inoculation and early PS19 tauopathy with altered sleep–wake organization and cortical EEG activity. The shared dark-phase increase in wakefulness is consistent with the established role of LC activity in promoting arousal, while the spectral changes are compatible with altered LC–noradrenergic modulation of cortical state [7, 28]. Whether LC hyperexcitability mediates these systems-level abnormalities remains to be determined.

### Retigabine attenuates bilateral LC tau pathology following unilateral fibril inoculation in PS19 mice

Tau fibril inoculation into the LC induces pathology in anatomically connected regions of PS19 mice [11], and increased neuronal activity can enhance tau release and propagation in experimental models [12]. Given the potassium-channel deficits and LC hyperexcitability identified above, we next asked whether chronic Kv7 modulation could modify pathological outcomes following local tau seeding.

Tau fibrils were inoculated unilaterally into the right LC of PS19 mice at two months of age. Beginning at 2.5 months, mice received vehicle or the Kv7 channel opener retigabine (10 mg/kg, intraperitoneally) every two days until tissue collection at four months (Fig. 7A). AT8-positive cells and puncta were detected on both sides of the brainstem, including the ipsilateral and contralateral LC regions (Fig. 7B–C). Section-level comparisons showed that greater LC AT8- positive cell density and hemispheric AT8-positive area fraction, together with lower LC TH- positive cell density, ipsilaterally than contralaterally Fig. S9A–C). Mouse-level comparisons showed similar changes or trends of changes (Fig. S9L-N). These results support the efficiency of locally inducing tauopathy by unilateral fibril inoculation into PS19 mice.

**FIGURE 7.**
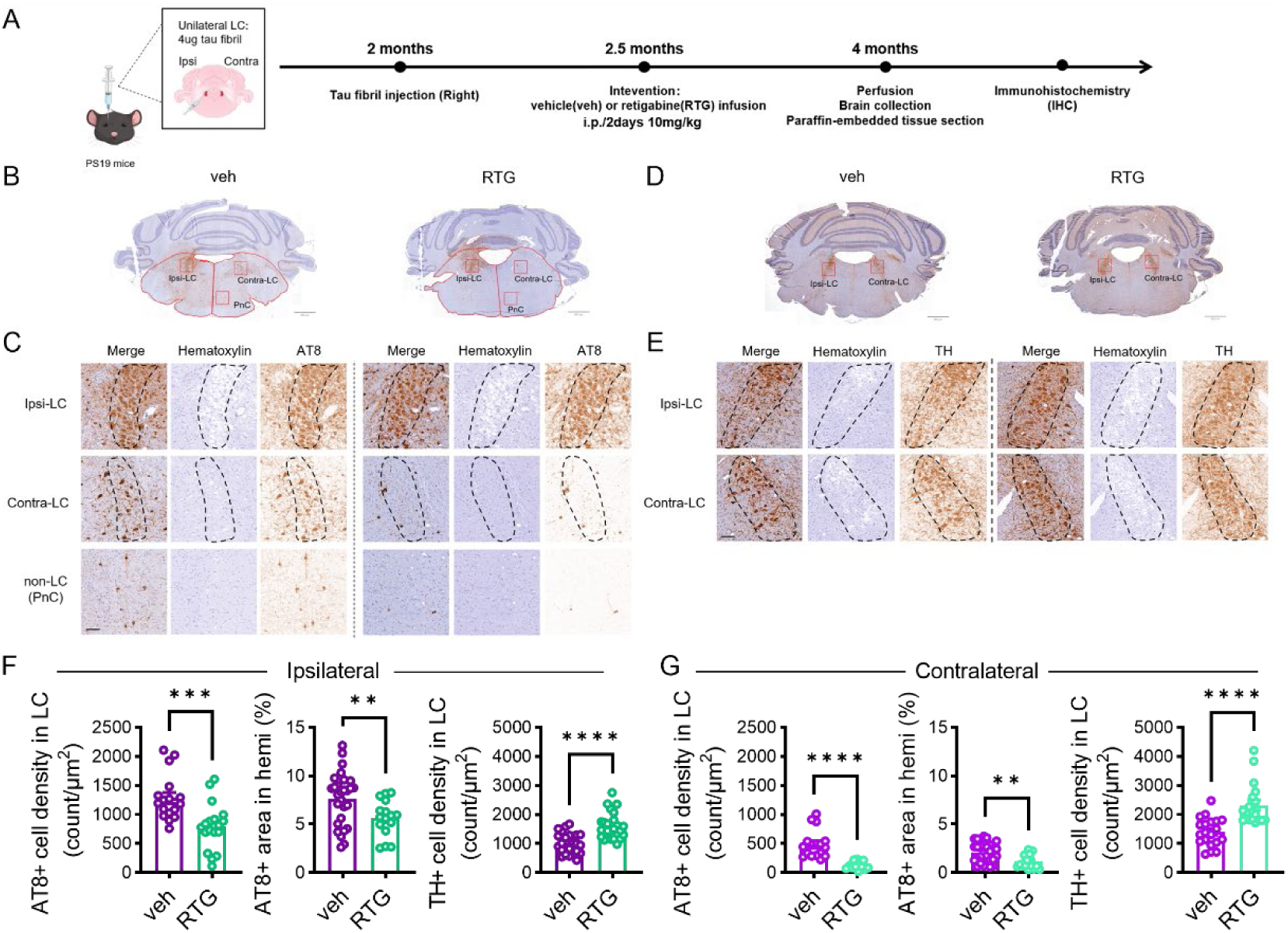
Retigabine rescues tau pathology, contralateral spreading, and LC neuronal loss after unilateral tau fibril injection into PS19 mice. (A) Experimental timeline. At 2 months of age, 4 μg of tau fibrils was injected unilaterally into the right LC of PS19 mice. Starting at 2.5 months, mice received vehicle (veh) or retigabine (RTG; 10 mg/kg, intraperitoneally, once every 2 days) until tissue collection at 4 months of age. (B-C) Representative sections from vehicle- and RTG-treated mice showing the regions examined for AT8 immunostaining. (B) Coronal brain slices. (C) Boxes indicate the ipsilateral LC, contralateral LC, and non-LC pontine nucleus (PnC) regions. Merged images, hematoxylin (purple), and AT8 (brown) are shown. Dashed outlines indicate the LC regions. (B: Scale bar, 800 μm; C: Scale bar, 50 μm). (D-E) Representative sections from vehicle- and RTG-treated mice showing the regions examined for TH immunostaining. (D) Coronal brain slices. (E) Boxes indicate the ipsilateral LC and contralateral LC. Merged images, hematoxylin (purple), and TH (brown) are shown. Dashed outlines indicate the LC regions. (D: Scale bar, 800 μm; E: Scale bar, 50 μm). (F-G) Quantification in the ipsilateral (F) and contralateral (G) brain regions. (veh: n = 3-4 mice; RTG: n = 3-4 mice, each mouse per 3-5 brain slices) Left panels: AT8-positive cell density in LC; Middle panels: AT8+ area in hemisphere (%); Right panels: TH+ cell density in LC. (F: left, p = 0.0006; middle, p = 0.0115; right, p < 0.0001; G: left, p < 0.0001; middle, p = 0.0026; right, p < 0.0001) Data are presented as [mean ± SEM] in Table S8. Mann–Whitney tests were performed.

Then we compared mice treated with retigabine to those with vehicle. Retigabine reduced AT8- positive cell density in the ipsilateral LC and the percentage of AT8-positive area in the corresponding brainstem hemisphere, while increasing ipsilateral LC TH+ cell density (Fig. 7B– F). The same pattern was observed contralaterally: retigabine reduced LC AT8-positive cell density and hemispheric AT8-positive area percentage and increased LC TH+ cell density (Fig. 7G). LC ROI areas did not differ significantly between treatment groups (Fig. S9E–H). The contralateral-to-ipsilateral ratio of AT8-positive area percentage was also lower after treatment (Fig. S9I). When measurements were averaged within each mouse, the treatment comparisons showed the same directional changes (Fig. S9J, K, O).

Together, these results show that chronic retigabine treatment initiated after unilateral LC fibril inoculation may attenuate bilateral AT8-defined tau pathology and is associated with higher TH+ cell density. These findings propose Kv7 modulation may serve as a pharmacological approach to modifying pathological outcomes in this model.

## DISCUSSION

Our findings identify early LC hyperactivity and hyperexcitability in two tauopathy models, accompanied by reduced low-threshold potassium currents. These abnormalities coincided with stress-associated transcriptional changes, reduced expression of neuronal-maintenance and potassium-channel genes, and increased nuclear REST. Both models exhibited altered sleep–wake organization and cortical oscillatory activity. Pharmacological modulation of potassium channels or GSK-3β inhibition attenuated LC hyperexcitability, while retigabine reduced bilateral LC AT8 burden and was associated with higher TH-positive neuronal density after unilateral fibril inoculation in young PS19 mice. Together, these findings implicate potassium-current dysfunction in early tau-associated LC abnormalities and support excitability as a potential target for modifying subsequent pathology.

Early LC dysfunction is increasingly evident across models of AD-related proteinopathy. Our recent study identified increased tonic firing and intrinsic excitability in young 5xFAD mice, associated with impaired Kv4/Kv7 regulation and dark-phase sleep disruption [29]. The present findings extend this phenotype to young PS19 mice and wild-type mice following local tau fibril inoculation. Increased spontaneous LC firing has also been reported following LC-targeted pathogenic tau expression and in *App* knock-in mice, while TgF344-AD rats exhibit enhanced bursting and aspects of stimulus-evoked hyperactivity [16, 30–32]. These studies converge on early dysregulation of LC activity, with its precise expression varying across models, disease stages and recording conditions.

Human imaging findings support the relevance of early LC dysfunction. Combined fMRI and PET studies have linked greater resting LC functional network activity to higher tau burden in amyloid-positive, cognitively unimpaired older adults [33]. Conversely, lower novelty-related LC function and connectivity have been associated with amyloid-related cognitive decline and subsequent cortical atrophy [34, 35]. These observations suggest disrupted regulation of LC function across behavioral states. Our age comparison likewise suggests stage-dependent abnormalities: young PS19 neurons were hyperactive, whereas surviving neurons sampled at 10 months showed firing rates comparable to controls. Reduced tonic and evoked activity has also been reported in aged P301S mice [36]. Collectively, these findings suggest that early LC dysfunction warrants investigation as an intervention target.

The sleep phenotypes provide a functional context for these activity changes. Both models examined here showed increased wakefulness and reduced NREM sleep during the dark phase, resembling the early hyperarousal phenotype in 5xFAD mice[29]. PS19 mice additionally showed reduced wakefulness and more fragmented wake bouts during the light phase, potentially reflecting the broader effects of brain-wide mutant tau expression. In comparison, LC-targeted P364S tau expression reduced NREM spindle density and duration while largely preserving sleep macrostructure[37]. This diversity is consistent with the multiple roles of LC-derived norepinephrine in regulating arousal and sleep microarchitecture[7, 38]. LC dysfunction may therefore contribute to shared early sleep abnormalities, with the resulting phenotype shaped by activity patterns and involvement of additional circuits.

At the cellular level, amyloidogenic and tauopathy models appear to converge on impaired potassium-current regulation through different upstream mechanisms. In young 5xFAD mice, soluble amyloid-β disrupted α2A-adrenergic receptor regulation of Kv4/Kv7 currents without detectable reductions in the measured channel transcripts or total protein abundance[29]. In young PS19 mice, potassium-current deficits accompanied reduced *Kcnq3* and *Kcnd1* expression and lower Kv7.3 and Kv7.2 immunoreactivity. Thus, early channel dysfunction may involve altered receptor-dependent regulation or reduced channel abundance. Reduced Kv7.2 protein despite no corresponding decrease detected in the exploratory transcriptomic dataset further suggests post-transcriptional regulation, with potential contributions from protein turnover, post-translational modification, trafficking and channel assembly.

These convergent deficits suggest an interaction between amyloid, neuronal activity and tau. Longitudinal human studies indicate that amyloid accumulation can precede detectable cortical fibrillar tau [39]. Experimental studies further show that amyloid plaques facilitate tau-seeded pathology, while endogenous tau contributes to amyloid-induced neuronal dysfunction [40, 41]. Our current and previous findings suggest a possible LC-centered feedback mechanism: soluble amyloid could impair receptor-dependent potassium-channel regulation leading to increased LC activity. Neuronal hyperexcitation alters tau phosphorylation, while increased activity enhances tau release and propagation [12, 42]. Subsequent tau-associated reductions in channel expression could reinforce the physiological deficit. This progression from functional channel dysregulation to reduced channel abundance provides a testable framework linking amyloid and tau to LC dysfunction.

REST provides a candidate link between early tau-associated stress and reduced Kv7 expression. REST can repress *Kcnq2* and *Kcnq3*, reducing M-current density and increasing neuronal excitability in other neuronal systems[25]. Increased nuclear REST in PS19 LC neurons therefore offers a plausible explanation for the accompanying reduction in *Kcnq3*. Importantly, REST also supports neuronal stress resistance during aging, and its induction at the onset of AD-related pathology can protect against amyloid and tau accumulation[26, 43]. Its elevation may represent an adaptive stress response with distinct consequences for potassium-channel regulation. Determining whether REST mediates the channel deficit, and whether selective regulation of its channel targets preserves its protective functions, will be important next steps.

The broader transcriptional changes suggest that early LC dysfunction extends beyond ion- channel regulation. Increased expression of genes such as *Mt1*, *Cst3* and *Apoe* is consistent with engagement of antioxidant, protease-regulatory and lipid-handling processes. These functions could support adaptation to cellular stress: cystatin C can promote protective autophagy, while apolipoprotein-mediated lipid transfer contributes to neuron–glia responses to oxidative stress[44, 45]. Conversely, the predominance of downregulated genes, including neurofilament genes and potassium-channel subunits, suggests reduced expression of programs associated with neuronal structure and maintenance. The coexistence of stress- response induction and declining maintenance-associated expression may represent an early state of incomplete compensation that leaves neurons vulnerable to subsequent challenges. These molecular programs are particularly relevant to selective LC vulnerability. Human comparisons between the LC and substantia nigra have identified differences in antioxidant pathways and cholesterol homeostasis[46]. A recent spatial transcriptomic study further links human LC molecular variation to APOE genotype, neuromelanin and local astrocytes[47]. Our findings complement these observations by connecting early tauopathy with physiological dysfunction and candidate transcriptional changes in marker-defined noradrenergic cells. Establishing which changes are adaptive, which impair neuronal function and which predict later degeneration will clarify their contribution to LC vulnerability.

The retigabine findings provide a rationale for targeting excitability during this early phase. Increased neuronal activity enhances tau release and propagation, and prolonged LC stimulation worsens tau phosphorylation and associated abnormalities in PS19 mice[12, 48]. In our unilateral seeding paradigm, retigabine reduced bilateral LC AT8 burden, consistent with modifying an activity-sensitive component of tau pathology. Higher TH-positive cell density additionally suggests preservation of the noradrenergic phenotype. Because treatment was systemic, LC-restricted interventions will be important for establishing the contribution of local Kv7 modulation.

Restoring appropriate LC activity could also influence sleep and broader noradrenergic functions. Local LC administration of guanfacine, an α2A-adrenergic receptor agonist, or retigabine improved sleep–wake disturbances in our previous 5xFAD study, supporting the feasibility of targeting this pathway[29]. Extending this approach to tauopathy may be relevant because sleep deprivation increases extracellular tau and enhances experimental tau spreading [13]. Norepinephrine also regulates microglial inflammatory responses and amyloid clearance, and loss of noradrenergic innervation can disrupt vascular tight-junction organization [49, 50]. These functions favor interventions that restore physiological LC activity while preserving beneficial noradrenergic signaling.

Together, our findings implicate potassium-current dysfunction in early tau-associated LC hyperexcitability. Convergence with amyloidogenic models identifies LC excitability as a shared point of physiological vulnerability and supports selective channel modulation as a potential strategy for modulating LC function and limiting subsequent pathology.

This study used male mice only, and the transcriptomic analysis comprised one pooled library per genotype. The expression findings are therefore exploratory, and their generalizability across sexes remains unresolved.

## Acknowledgement

We thank the staff members of the animal facility at the National Facility for Protein Science in Shanghai (NFPS), Shanghai Advanced Research Institute, Chinese Academy of Sciences, China for excellent support. This work was supported by Lin Gang Laboratory (Grant No. LGADB410101), Shanghai Municipal Science and Technology Major Project (Grant No. 2019SHZDZX02) and partly by the open funds of the State Key Laboratory of Brain Function and Disorders to Kai-Wen He.

## Author contributions

Conceptualization, K.-W.H.; Primary Investigation, Y.-C.Z., Y.-G.W., X.-T.Z.; Assistant Investigation, P.-Y.C., K.T., Z.-Y.Z., X.M.; Material preparation: W. Z.; Z.-H.H.; Data Analysis, Y.- C.Z., Y.-G.W., X.-T.Z., Z.-Y.Z.; P.-Y.C; Writing, Y.-C.Z., X.-T.Z., and K.-W.H.; Funding Acquisition, H.-T.X., K.-W.H; Supervision, K.-W.H.

## METHODS

### Animals

All procedures were approved by the Institutional Animal Care and Use Committee of the Interdisciplinary Research Center on Biology and Chemistry, Chinese Academy of Sciences. Mice were group-housed under a 12-h light/12-h dark cycle with food and water available ad libitum. Animals included male C57BL/6J mice purchased from SPF (Beijing) Biotechnology, PS19 mice and their wild-type littermates, and tau-knockout mice (B6.129X1-Mapt^tm1Hnd^/J, originally obtained from The Jackson Laboratory, referred to as mTauKO). Purchased C57BL/6J mice and PS19 wild-type littermates served as controls for the fibril-inoculation and transgenic experiments, respectively.

### Electroencephalographic and electromyographic recordings

Electroencephalographic (EEG) and electromyographic (EMG) recordings were performed as previously described [51]. Briefly, under isoflurane anesthesia (1–2%), mice were secured in a stereotaxic frame, and the skull was exposed. Two epidural screw electrodes were implanted at the reported coordinates relative to bregma: anteroposterior, +1.5 to −3 mm; mediolateral, +1.5 mm. Two insulated stainless-steel wires with exposed tips were implanted in the left and right dorsal neck muscles for EMG recording. Electrodes were connected to a four-pin socket secured to the skull with dental cement. The incision was sutured and treated with triple-antibiotic ointment. Mice recovered for at least 10 days.

Mice were habituated to the recording chamber for 24 h before continuous EEG/EMG recording for 48–72 h. The second 24-h recording period was analyzed. The regular light/dark cycle and unrestricted access to food and water were maintained. Signals were acquired using Spike Hound at 1,024 Hz with a preamplifier gain of 1,000. The reported acquisition band-pass settings were 0.3–1,000 Hz for EEG and 1–5,000 Hz for EMG.

### Ex vivo slice electrophysiology

#### Acute brain-slice preparation

Acute brain slices for electrophysiology were prepared as described previously [29, 52] 1–2 h before light onset, corresponding to zeitgeber time (ZT) 22–23. Briefly, mice were anesthetized with isoflurane vapor and transcardially perfused with about 10 ml of ice-cold dissection buffer containing the following: 93 mM NMDG, 2.5 mM KCl, 1.2 mM NaH_2_PO_4_·2H_2_O, 30 mM NaHCO_3_, 20 mM HEPES, 5 mM sodium ascorbate, 2 mM Thiourea, 3 mM sodium pyruvate, 25 mM D-Glucose, 12 mM NAc, 10 mM MgCl_2_, 0.5 mM CaCl_2_ bubbled with 95% O_2_/ 5% CO_2_. Mice were then decapitated and whole brains were rapidly removed. Coronal slices containing the LC were cut at 300 μm using a vibratome (VT1200S, Leica) in ice-cold dissection solution. Slices were incubated in dissection solution at 30 °C for 15 min, then transferred to artificial cerebrospinal fluid (ACSF) containing, in mM: 119 mM NaCl, 5 mM KCl, 1.25 mM NaH_2_PO_4_, 1 mM MgCl_2_, 2 mM CaCl_2_, 26 mM NaHCO_3_, and 10 mM Dextrose, saturated with 95% O_2_ and 5% CO_2_). Slices recovered at room temperature for at least 1 h before recording.

#### Whole-cell electrophysiology

For whole cell recordings, slices were transferred to a recording chamber continuously perfused with 30 ± 0.5°C ACSF at 2 ml/min. Putative LC neurons were identified by anatomical location and morphology using an upright fluorescence microscope (XT640-W, Olympus). Borosilicate glass pipette recording electrodes had resistances of 3-6 MΩ. Unless otherwise specified, pipettes contained a potassium-based internal solution comprising, in mM: 130 mM K-gluconate, 10 mM KCl, 10 mM HEPES, 0.2 mM EGTA, 0.5 mM Na_3_GTP, 4 mM MgATP, 10 mM Na-phosphocreatine, adjusted to pH 7.2-7.4 and 285-300 mOsm. Cells were included when membrane resistance exceeded 100 MΩ and access resistance was below 25 MΩ; recordings were excluded if either value changed by more than 25%. Signals were low-pass filtered at 2 kHz and digitized at 10 kHz using Clampex (Axon).

##### a.#Spontaneous firing and intrinsic excitability

Spontaneous firing was recorded in current-clamp mode without injected current for at least 30 s. Cells with resting membrane potentials below −40 mV were included.

Intrinsic excitability was assessed in drug-free ACSF as previously described [53, 54]. Cells were maintained at −65 mV by current injection. Action-potential threshold and rheobase were estimated using a 500-ms current ramp from 0 to 400 pA. Current-evoked firing was assessed using 500-ms steps from 0 to 1,000 pA in 100-pA increments. Maximal firing and the current producing maximal firing (I_maxAP_) were determined from these responses. Cells with resting membrane potentials below −45 mV were included.

##### b.#Synaptic currents

Spontaneous excitatory postsynaptic currents (sEPSCs) were recorded at −60 mV in drug-free ACSF using the potassium-based internal solution [53, 54].

Miniature inhibitory postsynaptic currents (mIPSCs) were recorded at −70 mV in ACSF containing 1 μM tetrodotoxin, 100 μM DL-2-amino-5-phosphonovaleric acid (DL-APV) and 20 μM 6-cyano-7-nitroquinoxaline-2,3-dione (CNQX). The internal solution contained, in mM: 130 CsCl, 8 NaCl, 0.2 CaCl₂, 10 HEPES, 2 EGTA, 0.5 Na₃GTP, 4 MgATP and 5 QX-314, adjusted to pH 7.2–7.3 with CsOH and 285–300 mOsm.

##### c.#Voltage-gated potassium-current recordings

Potassium currents were recorded in voltage-clamp mode using the potassium-based internal solution [20]. The following drugs were added to the perfused ACSF to isolate Kv currents: 1 μM TTX, 100 μM DL-APV, 20 μM gabazine and 20 μM CNQX. Total outward potassium currents were recorded following a 750-ms conditioning step to −120 mV, followed by 400-ms test steps from −100 to +90 mV in 10-mV increments. Currents remaining after a 750-ms conditioning step to −30 mV were designated the high-threshold component. The low-threshold component was calculated by subtracting these currents from those recorded after conditioning at −120 mV.

The early component was quantified as the maximal current within the first 15 ms of the test step, and the persistent component as the plateau current at the end of the step. Current density was calculated by dividing current amplitude by cell capacitance.

##### d.#Pharmacological treatments

Retigabine (ezogabine; 20 μM) and NS5806 (10 μM) were bath-applied in ACSF to examine the effects of potassium-channel modulation on intrinsic excitability and spontaneous firing. The glycogen synthase kinase-3 inhibitor AR-A014418 (5 μM) was pre-incubated and bath-applied to assess its effects on potassium currents, intrinsic excitability and spontaneous firing.

#### Recombinant tau purification and fibril preparation

Full-length human tau (2N4R; 441 amino acids) carrying the P301L mutation was cloned into pRK172 and expressed in BL21-CodonPlus (DE3) RIPL cells (EC1077S, Weidibio). Tau was purified by cation-exchange fast protein liquid chromatography [55], followed by size-exclusion chromatography.

Bacteria were grown in antibiotic-supplemented LB medium at 37 °C. At an optical density of 1.0 at 600 nm, expression was induced with 1 mM isopropyl β-D-1-thiogalactopyranoside for 2 h. Bacteria were collected by centrifugation, resuspended in lysis buffer containing 50 mM Tris-HCl, 2 mM EGTA, 2 mM dithiothreitol and 2 mM phenylmethylsulfonyl fluoride (pH 8.0), and disrupted by high-pressure homogenization. Lysates were heated at 100 °C for 20 min and ultracentrifuged. The supernatant was dialyzed overnight at 4 °C against buffer containing 20 mM PIPES, 2 mM dithiothreitol and 2 mM phenylmethylsulfonyl fluoride (pH 6.5).

Cation-exchange and size-exclusion chromatography were performed using an ÄKTA system (GE Healthcare). Purified tau was dialyzed into 100 mM sodium acetate (pH 7.0), and protein concentration was measured using a bicinchoninic acid assay (E112-02, Vazyme).

Tau preformed fibrils were generated as previously described [35]. Recombinant tau (40 μM) was incubated with heparin sodium (40 μg/mL; A16198, Alfa Aesar), 2 mM dithiothreitol and 0.02% sodium azide in 100 mM sodium acetate (pH 7.0). Samples were agitated at 1,000 rpm and 37 °C for 5–7 days in a thermomixer (Eppendorf). Fibril formation was verified by negative- stain electron microscopy.

Fibrils were pelleted at 100,000 × g for 1 h, resuspended in phosphate-buffered saline (PBS), and pelleted again to remove soluble components. The final pellet was resuspended in PBS. Fibril concentration was estimated by SDS–PAGE, Coomassie staining and densitometry against monomeric tau standards. Fibrils were fragmented using a probe sonicator before use.

#### Primary hippocampal neuron cultures

Hippocampi were dissected from embryonic day 16–18 C57BL/6N embryos, dissociated and plated on poly-L-lysine-coated dishes. Cells were initially maintained in Dulbecco’s modified

Eagle medium, which was replaced within 2 h by Neurobasal-A medium supplemented with B27 (Invitrogen). Cultures were maintained at 37 °C in 5% CO₂. Monomeric tau or fibrils (1 μg) were added at 7 days in vitro, and neurons were collected for immunostaining or western blotting at 14 days in vitro.

#### Stereotaxic tau inoculation and retigabine treatment

Stereotaxic injections were performed as previously described [56]. 2-month-old wild-type mice were anesthetized with isoflurane vapor (1%–2%) and head-fixed in a stereotaxic frame. Bilateral injections into the LC were performed using a Hamilton syringe fitted with a glass electrode at a rate of 0.1 μL/min. The injection coordinates were as follows: Bregma, −5.45 mm; Lateral, ± 0.89 mm; Depth, −2.8 mm.

Wild-type mice were randomly assigned to receive Hep-T40PL preformed fibrils (fibril; 0.5 μg/μL), T40PL monomer (monomer, 0.5 μg/μL), or sterile D-PBS. A total of 2 μL of the assigned solution was injected into each site. PS19 mice received a unilateral injection of Hep-T40PL PFF (fibril, 2 μL, 2 μg/μL) into the right LC using the same procedure. After surgery, mice were returned to their home cages and monitored daily for one week. Unilaterally inoculated PS19 mice received retigabine (10 mg/kg, intraperitoneally) every other day until 4 months of age.

#### Tissue collection and immunoblotting

Brain tissue was collected 1–2 h before light onset. Mice were anesthetized with isoflurane, and acute slices containing the target regions were prepared in ice-cold NMDG dissection solution. After recovery in ACSF and incubation with AR-A014418 or vehicle, slices were rapidly frozen in liquid nitrogen and stored at −80 °C.

Cultured neurons were placed on ice, lysed and gently agitated for 10 min. Tissue and cell samples were homogenized in radioimmunoprecipitation assay buffer containing 50 mM Tris- HCl (pH 7.4), 1% NP-40, 150 mM NaCl, 0.25% sodium deoxycholate, 1 mM EDTA, 1 mM sodium orthovanadate, 1 mM NaF and 1 mM phenylmethylsulfonyl fluoride, supplemented with protease and phosphatase inhibitor cocktails (B14002 and B15002, Bitool). Tissue was homogenized using a grinder (Z190179, Sigma).

Lysates were centrifuged at 20,000 × g for 20 min at 4 °C. Supernatants were mixed with 4× loading buffer to approximately 1 μg/μL and heated at 95 °C for 5 min. Protein samples (10–20 μg) were separated on 10% SDS–polyacrylamide gels and transferred to polyvinylidene difluoride membranes (Bio-Rad). Membranes were blocked in Tris-buffered saline containing Tween-20 and 5% nonfat milk for 2 h at room temperature, then incubated overnight at 4 °C with AT8 (1:500), tau (1:40,000), actin (1:1,000), or tubulin (1:40,000) antibodies. After washing, membranes were incubated with secondary antibodies for 1–2 h at room temperature. Chemiluminescence was acquired and analyzed using a Fusion imaging system (VILBER). Antibody details are provided in Table S1.

### Histology and immunofluorescence

#### Tissue preparation and immunohistochemistry

Mice were anesthetized with isoflurane vapor and transcardially perfused with 10 ml PBS followed by 10 ml of 10% neutral buffered Formalin (NBF) for immunohistochemistry or 4% phosphate buffered paraformaldehyde (PFA) for immunofluorescence. Brains were then removed and fixed overnight in NBF or PFA.

Formalin-fixed brains were rinsed in PBS, embedded in paraffin and sectioned coronally at 6 μm using a microtome (RM2125 RTS, Leica). Sections were deparaffinized, rehydrated through graded ethanol (100%, 95%, 80% and 70%), and blocked in 0.1 M Tris containing 2% fetal bovine serum for 1 h at room temperature. Sections were incubated overnight at 4 °C with AT8 (1:2,000) or tyrosine hydroxylase (TH; 1:1,000) antibodies, followed by horseradish peroxidase- conjugated secondary antibodies. Staining was developed using 3,3′-diaminobenzidine (DAB) and counterstained with Mayer’s hematoxylin. All antibodies and materials used are listed in Table S1.

#### Immunofluorescence staining (IF)

PFA-fixed brains were dehydrated in 20% phosphate-buffered sucrose for 1 day and 30% phosphate-buffered sucrose for 2 days before embedded in tissue freezing medium (#14020108926, Leica) and rapidly frozen in liquid nitrogen. 20 μm-thick coronal brain slides containing LC or hippocampus were section by Cryostat (CM3050 S, Leica).

Sections underwent antigen retrieval using P0090 reagent (Beyotime), followed by blocking for 1 h at room temperature in PBS containing 5% normal goat serum, 1% IgG-free bovine serum albumin and 0.1% Triton X-100. Sections were incubated overnight at 4 °C with AT8 (1:500), TH (1:3,000), NeuN (1:2,000), Kv7.2 (1:500), Kv7.3 (1:500), GFAP (1:10,000), Iba1 (1:1,000), or REST (1:500) antibodies. After washing in PBS containing Tween-20, sections were incubated with secondary antibodies for 2 h at room temperature, washed and mounted. Nuclei were stained with 4′,6-diamidino-2-phenylindole for 10 min, and sections were coverslipped with ProLong Gold Antifade Mountant (P36930, Invitrogen). Antibody and reagent details are provided in Table S1.

#### Imaging

All fluorescence signal was acquired using a spinning disk microscope (Dragonfly 200, Andor) connected to camera (Zyla sCMOS, Andor) with a 20x air lens. The image resolution is x/y: 0.300903/ 0.301040 μm/pixel. Stack images were acquired for all images. Step size (μm) and thickness (μm) of were as followed: 0.5, 1.

Immunohistochemical staining signal was acquired using P250 slide scanner (3DHISTECH), with 20x air lens. The image resolution is x/y: 0.2428/ 0.2428 μm/pixel. Single plane images were acquired.

#### Single-cell RNA sequencing

Mice were anesthetized with isoflurane and transcardially perfused with approximately 10 mL of ice-cold, carbogenated NMDG dissection solution. Coronal slices containing the LC were prepared at 300 μm using a vibratome (VT1200S, Leica). LC-containing tissue was collected by punch. Tissue from eight mice of each genotype was pooled to generate one sample per genotype.

Pooled tissue was digested in oxygenated NMDG/HEPES solution containing papain (10 U/mL; LK003178, Worthington) and DNase I (1,000 U/mL; D4527-10KU, Sigma) at 37 °C with gentle shaking for 30 min. Tissue was centrifuged at 400 × g for 5 min at 4 °C and resuspended in 2 mL of oxygenated NMDG/HEPES solution containing 2% fetal bovine serum (10099-141, Gibco) and DNase I (10 U/mL).

Tissue was sequentially triturated through glass pipettes with 300- and 200-μm tip diameters until no visible clumps remained. Cells were collected at 400 × g for 5 min at 4 °C. For debris removal, the pellet was resuspended in 1,550 μL NMDG solution and mixed with 450 μL debris- removal solution (130-109-398, Miltenyi). After overlaying 2 mL NMDG solution, samples were centrifuged at 3,000 × g for 10 min with mild acceleration and deceleration. The bottom fraction was collected.

Suspensions were passed through a pre-rinsed 40-μm strainer (352235, Falcon), diluted to 10 mL with resuspension solution, and centrifuged at 400 × g for 5 min at 4 °C. Cells were resuspended in 50 μL oxygenated NMDG/HEPES solution containing 2% fetal bovine serum. Cell concentration and viability were assessed by trypan blue exclusion using a hemocytometer. Libraries were prepared using the DNBelab C Series High-throughput Single-cell RNA Library Preparation Kit V3.0 (MGI Tech) according to the manufacturer’s instructions and sequenced on a DNBSEQ-T7 platform to approximately 500 million reads per sample.

## QUANTIFICATION AND STATISTICAL ANALYSIS

### Sleep and EEG analysis

Wakefulness, NREM sleep and REM sleep were automatically classified in 5-s epochs using NeuroScore (Data Sciences International), with a slow-wave ratio threshold of 0.4 and theta ratio threshold of 3. The results were manually checked. The wake epochs shorter than 10 sec were defined as microarousal.

EEG signals were band-pass filtered at 0.5-100 Hz and power line filtered at 50 Hz before analyzed by Fourier Transform. The brain oscillation bands were defined as Delta (δ): 0.5-4 Hz, Theta (θ): 4-8 Hz, Alpha (α): 8-12 Hz, Beta (β): 12-30 Hz, Gamma (γ): >30 Hz. The Unitary power was defined as average power per 5-second epoch. Each bout is defined as a continuous state of wake/ NREM/ REM with no interruptions for sleep structure analysis.

### Synaptic current analysis

sEPSCs and mIPSCs were analyzed using the MiniAnalysis program (Synaptosoft, Decatur, GA) as described previously [53]. Event detection threshold was set at 3 times over the RMS noise. At least 300 events with rise time < 3 msec were selected for each cell to calculate sEPSC and mIPSCs frequency and amplitude.

### Fluorescence-image analysis

Images were batch-analyzed using Fiji/ImageJ and Imaris (Oxford Instruments). Z-stacks were maximum-intensity projected and background-subtracted. LC neuronal regions of interest (ROIs) were defined by TH labeling; neuronal ROIs in other regions were defined by NeuN labeling. Somatic AT8 fluorescence was measured in the dentate gyrus, CA1, CA3, striatum, pedunculopontine nucleus, dorsal raphe, periaqueductal gray and laterodorsal tegmental nucleus.

For LC Kv7.2 and Kv7.3 measurements, somatic ROIs delineated on the TH channel were transferred to the corresponding channel to measure mean fluorescence intensity. REST fluorescence was measured in whole-cell and nuclear ROIs. Nuclear ROIs were defined by the intersection of each neuronal ROI with its DAPI-positive nuclear mask.

TH-positive LC profiles and Iba1-positive puncta were detected using the Imaris Spots module. Image analysis was performed blinded to experimental group. Mean AT8, TH, GFAP, REST, Kv7.2 and Kv7.3 intensities were normalized to the corresponding control group. When experiments were combined across batches, each batch was normalized to its own control. TH- positive cell and Iba1-positive puncta densities were reported without normalization.

### Immunohistochemical analysis

DAB-stained, hematoxylin-counterstained sections were analyzed using QuPath. LC ROIs were manually delineated in each hemisphere using cytoarchitecture, including the relatively large neuronal profiles of the LC. AT8- or TH-positive profiles with identifiable somata were manually counted within each ROI. Profile density was calculated as the count divided by ROI area and expressed as cells/mm².

For hemispheric AT8 burden, the brainstem tissue area in each hemisphere was manually delineated. AT8-positive staining was identified using an identical threshold across images. AT8-positive area fraction was calculated as the positive area divided by the analyzed hemispheric tissue area. ROI delineation and counting were performed blinded to experimental group.

### Single-cell RNA sequencing analysis

Gene–barcode count matrices were imported into R using DropletUtils. The initial matrix contained 34,285 features and 28,705 barcodes. Potential doublets were scored separately for each sample using the scds hybrid method, cxds_bcds_hybrid. The expected number of doublets was estimated as 0.01 × n²/1,000, where n was the number of barcodes in that sample; barcodes with the highest scores were removed accordingly.

The remaining counts were imported into Seurat. Genes detected in fewer than three cells were excluded. Cells were retained if they contained >500 and <9,000 detected genes and <5% mitochondrial transcripts.

For global analysis, counts were normalized using NormalizeData, and 3,000 highly variable genes were selected using the variance-stabilizing transformation method. Data were scaled and analyzed by principal component analysis. The first 14 principal components were used for nearest-neighbor graph construction, uniform manifold approximation and projection (UMAP), and Leiden clustering (algorithm 4; resolution 0.1).

Cell identities were assigned using canonical markers: *Snap25, Rbfox3, Syp, Tubb3* and *Map2* for neurons; *Slc17a7* and *Slc17a6* for excitatory neurons; *Gad1, Gad2, Slc6a1* and *Slc32a1* for inhibitory neurons; and *Th, Dbh, Slc6a2, Slc18a2, Ddc, Phox2a, Phox2b* and *Gch1* for noradrenergic neurons.

Neurons were subsetted and independently reprocessed using normalization, selection of 2,000 highly variable genes, scaling and principal component analysis. The first 13 principal components were used for UMAP, nearest-neighbor graph construction and Leiden clustering at resolution 0.5. Neuronal clusters were annotated from their marker-expression profiles.

For differential-expression analysis, marker-defined noradrenergic-like neurons were selected by detectable expression of at least one of *Th, Dbh, Slc6a2* or *Slc18a2*. PS19 and WT cells were compared using the Wilcoxon rank-sum test implemented in FindMarkers. P values were adjusted using the Benjamini–Hochberg procedure. Genes with adjusted P ≤ 0.05 and absolute log₂ fold change ≥0.25 were included in the displayed differential-expression summaries.

### Immediate-early-gene signature scoring

A rank-based enrichment score adapted from UCell [57] was calculated as neural activity score for the following 16-gene signature: *Fos, Fosb, Jun, Junb, Egr1, Egr2, Egr3, Egr4, Arc, Npas4, Nr4a1, Nr4a2, Nr4a3, Bdnf, Homer1* and *Rgs2*.

Within each cell, genes were ranked by descending normalized expression, with average ranks assigned to ties. Ranks greater than or equal to the maximum rank were capped at *R_max_*, defined as the smaller of 1,500 and the total number of genes in the expression matrix. For cell c and signature G containing n genes, the score was calculated as follows

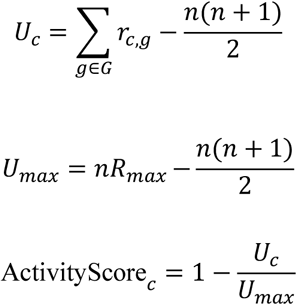

Here, *G* denotes the IEG gene set and *r_c_*_,*g*_ is the rank of gene g in cell c. Higher score indicate greater enrichment of the IEG signature.

### Statistical analysis

Sample size is indicated in all figures as mouse number or cell then mouse number. Statistical analysis was performed by Prism V6.0 software (GraphPad Software, Inc.). Wilcoxon signed- rank test was used for paired data. Mann-Whitney U test was used for unpaired data. Multiple t-test were used to analysis power spectrum in light/dark phase and absolute power comparison. Two-way *ANOVA* with Šídák multiple-comparison test was used to compare I-V curves, I-O curves and power ratio analysis. Pearson correlation coefficient was calculated for correlation analysis.

Data are presented as mean ± standard error of the mean. Statistical significance was defined as P < 0.05, with *P < 0.05, **P < 0.01, ***P < 0.001 and ****P < 0.0001. P values between 0.05 and 0.1 were displayed numerically.

## Code availability

Custom MATLAB code is available at: https://github.com/KaiWen-Helab/Zhang_tau2024

## SUPPLEMENTARY FIGURES AND LEGENDS

**FIGURE S1.**
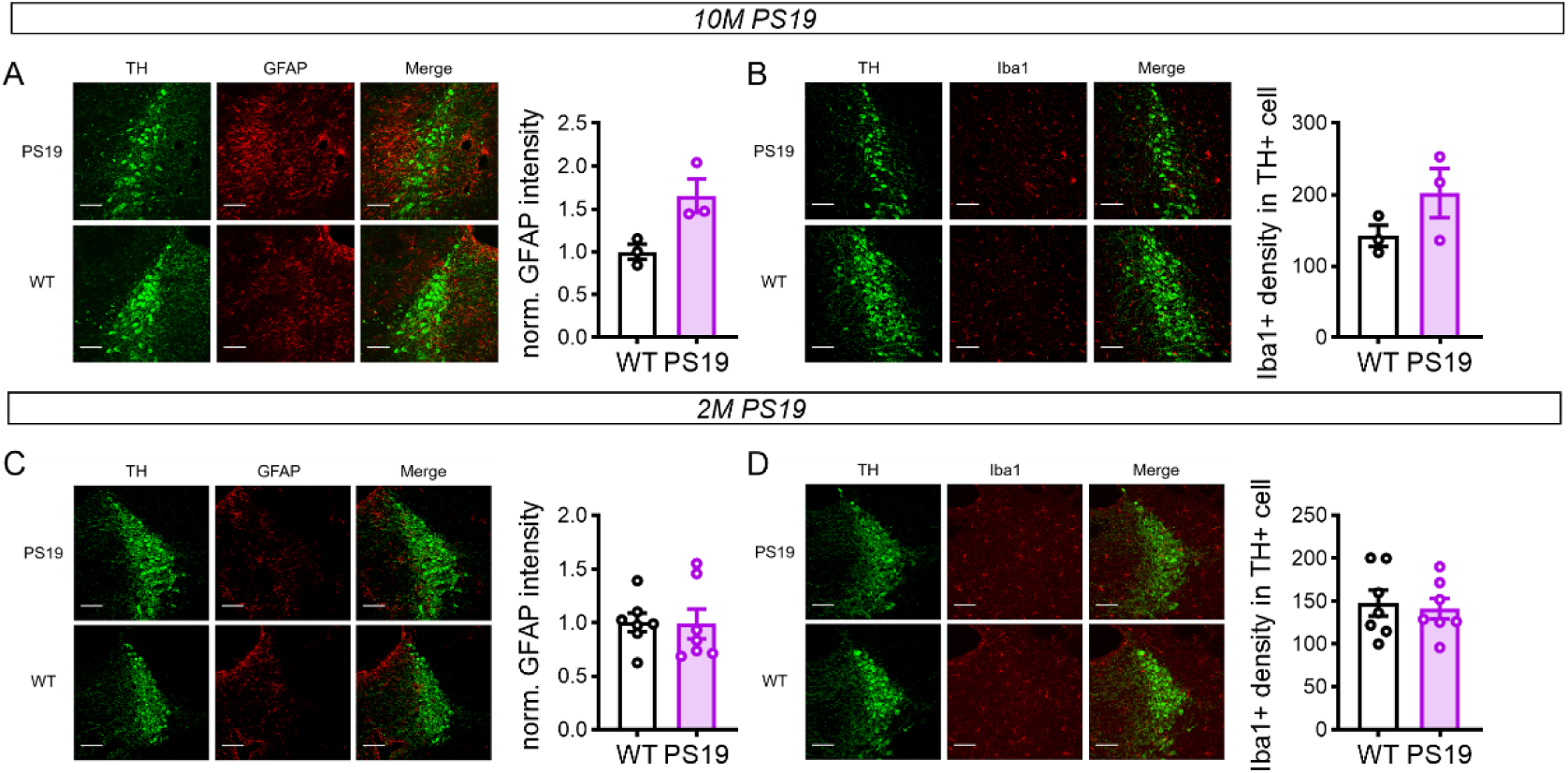
Neuroinflammation in 10-month-old and 2-month-old PS19 mice. (A-B) Immunostaining of the LC in 10-month-old WT and PS19 mice. (WT: n = 3 mice; PS19: n = 3 mice) (A) Representative LC images (left) showing tyrosine hydroxylase (TH, green), GFAP (red) and quantification of signals (right). (B) Representative LC images (left) showing tyrosine hydroxylase (TH, green), Iba1 (red) and quantification of density (right). (Scale bar, 100 μm) (C-D) Immunostaining of the LC in 2-month-old WT and PS19 mice. (WT: n = 7 mice; PS19: n = 7 mice) (C) Representative LC images (left) showing tyrosine hydroxylase (TH, green), GFAP (red) and quantification of signals (right). (D) Representative LC images (left) showing tyrosine hydroxylase (TH, green), Iba1 (red) and quantification of density (right). (Scale bar, 100 μm) Data are presented as [mean ± SEM] in Table S2. Mann–Whitney tests were performed.

**FIGURE S2.**
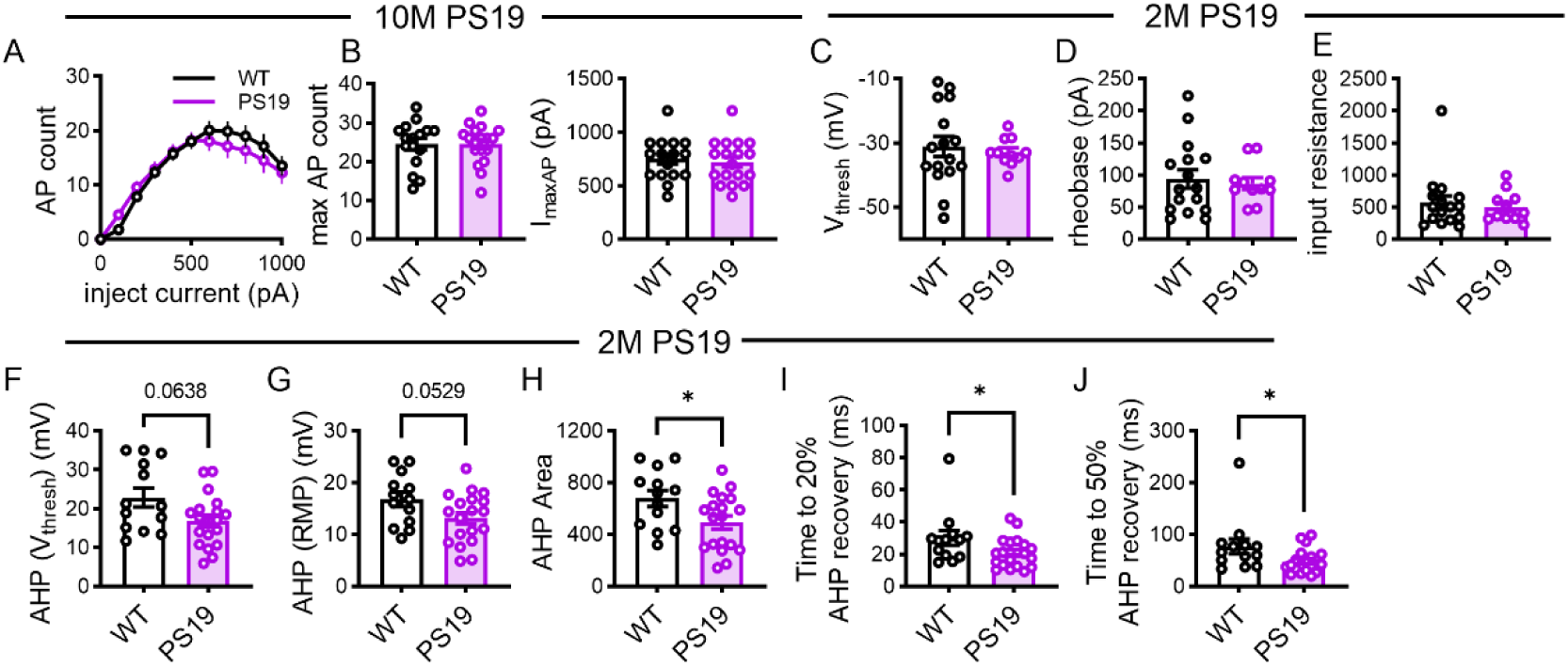
Electrophysiological properties of LC neurons in 2-month-old PS19 mice. (A-B) Comparison of neuronal excitability by injecting current steps in 10-month-old PS19 mice LC neurons (WT: 16 cells from 4 mice; PS19: 19 cells from 4 mice). (A) Current-spiking relationship of LC neurons. (B) Quantification of maximal firing rate (left) and currents required to generate the maximal firing (I_maxAP_, right). (C-E) Comparison of AP threshold (C), rheobase (D), and input resistance (E) in 2-month-old PS19 mice LC neurons (WT: 16 cells from 4 mice; PS19: 11 cells from 4 mice) (F-J) Comparison of afterhyperpolarization (AHP) potential measured relative to AP threshold (F) or resting membrane potential (G), respectively, AHP area (H), and time spent to recover 20% (I) or 50% AHP (J) in 2-month-old PS19/WT mice LC neurons (WT: 13 cells from 4 mice; PS19: 19 cells from 4 mice) (H: p = 0.0328; I: p = 0.0266; J: p = 0.0361). Data are presented as [mean ± SEM] in Table S3. Mann–Whitney tests were performed.

**FIGURE S3.**
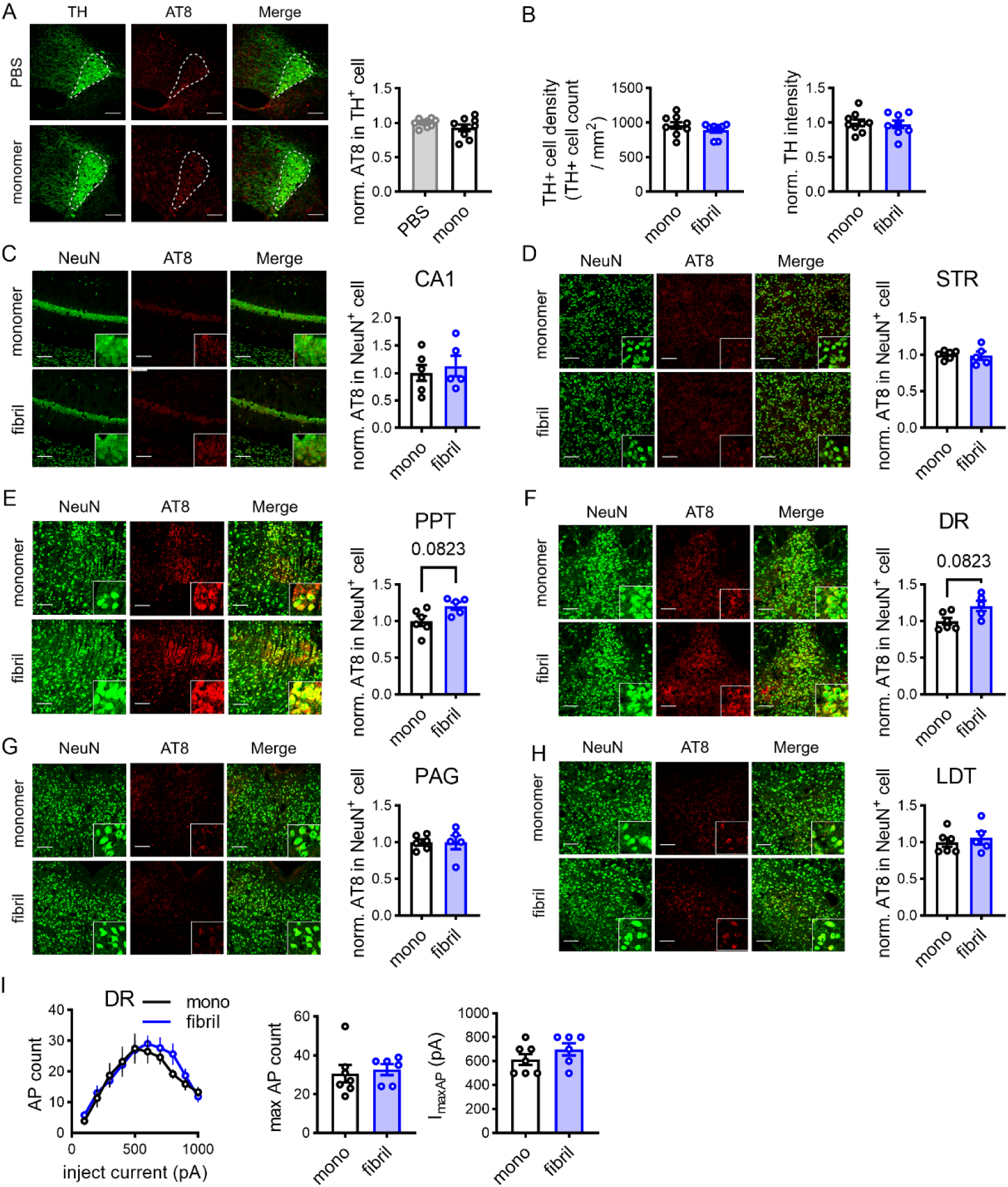
Tau fibril inoculation effect on various brain regions. (A) Immunostaining representative images (left) and quantification (right) of tau phosphorylation 1-month post tau monomer inoculation into LC. (red: AT8, green: TH, scale bar, 100 µm, LC, dotted line; PBS: n = 9 mice; mono: n = 9 mice) (B) Quantification of TH-positive LC neuronal density (left) and normalized TH intensity (right) (mono: n = 9 mice; fibril: n = 8 mice) (C–H) Immunostaining analysis of tau phosphorylation in various brain regions post fibril inoculation into LC. C: CA1, D: striatum, E: Pedunculopontine tegmental nucleus, F: Dorsal Raphe, G: Periaqueductal gray, H: Laterodorsal tegmental nucleus. Left panels, Representative images of AT8 signal in monomer- or fibril-inoculated wild-type mice (red: AT8; green, NeuN, scale bar, 100 µm). Right panels, Quantification of the normalized AT8 signal intensity per NeuN-positive soma. (mono: n = 6 mice; fibril: n = 5 mice) (I) Dorsal raphe neuronal excitability remained unaltered in wild-type mice inoculated with tau fibril into LC. Left, current-spiking relationship of neurons of LC-fibril- (blue) or LC-monomer- inoculated (black) wild-type mice. Middle, maximal firing rates. Right, I_maxAP_. (mono: 7 cells from 2 mice; fibril: 6 cells from 2 mice) Data are presented as [mean ± SEM] in Table S4. Mann–Whitney tests were performed.

**FIGURE S4.**
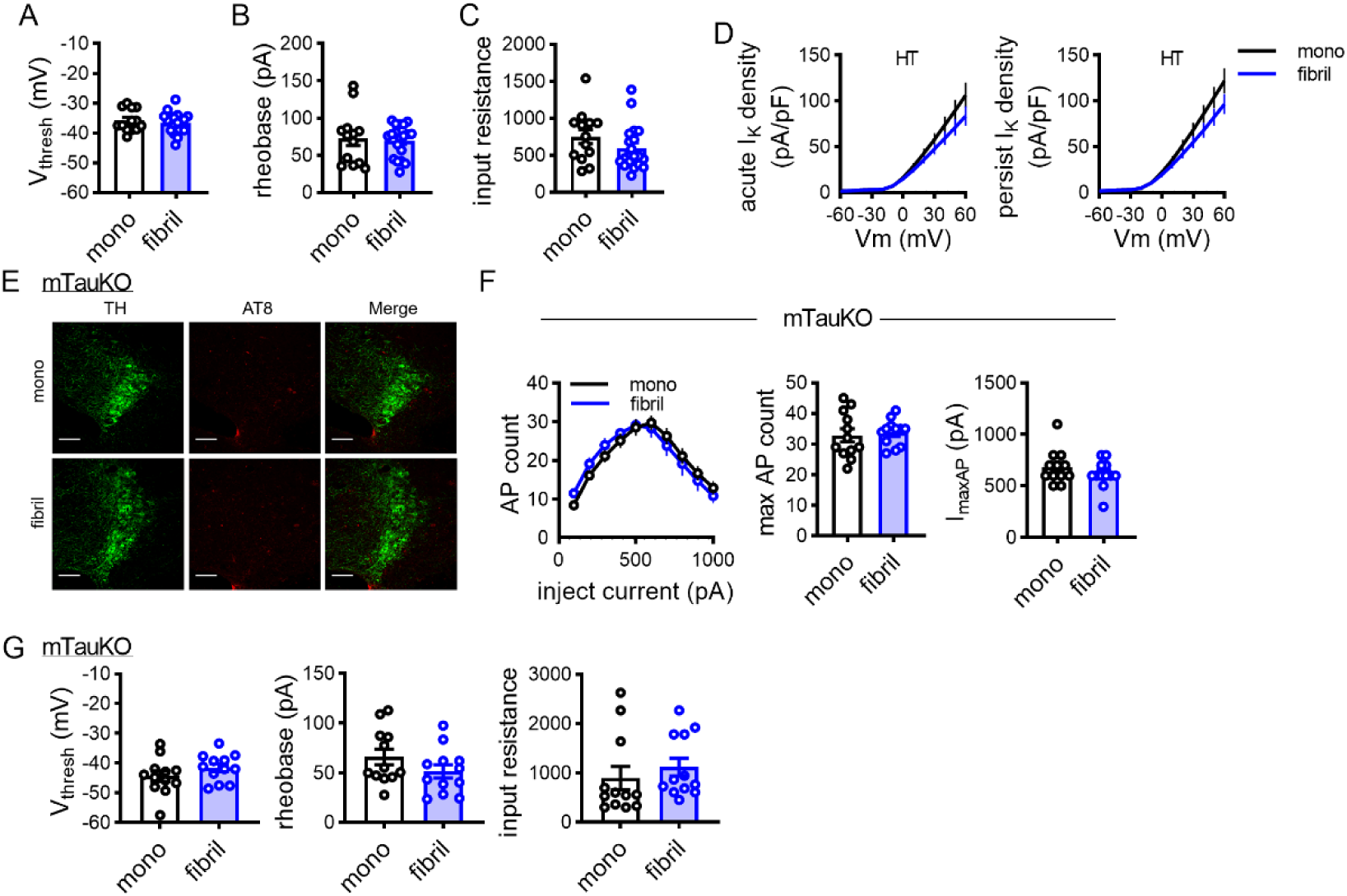
Electrophysiological properties of LC neurons in tau fibril-inoculated mice. (A-C) Comparison of action potential threshold potential (A), rheobase (B) and input resistance (C) of LC neurons of monomer- or fibril-inoculated wild-type mice. (mono: 13 cells from 3 mice; fibril: 19 cells from 4 mice) (D) V-I curves of high threshold (HT) I_k_ density. Left panels, acute Ik density; Right panels, persist Ik density. (mono: 12 cells from 3 mice; fibril: 13 cells from 3 mice) (E) Lack of AT8 signal in the mono- or fibril-inoculated mTauKO mice. (Scale bar, 100 µm) (F-G) Fibril failed to induce hyperexcitability in LC neurons of mTauKO mice. (F) Left, current- spiking relationship; Middle, quantification of maximal firing rate; Right, I_maxAP_. (G) Left, action potential threshold potential; Middle, rheobase; Right, input resistance. (mono:12 cells from 3 mice; fibril: 12 cells from 3 mice) Data are presented as [mean ± SEM] in Table S4. Mann–Whitney tests were performed.

**FIGURE S5.**
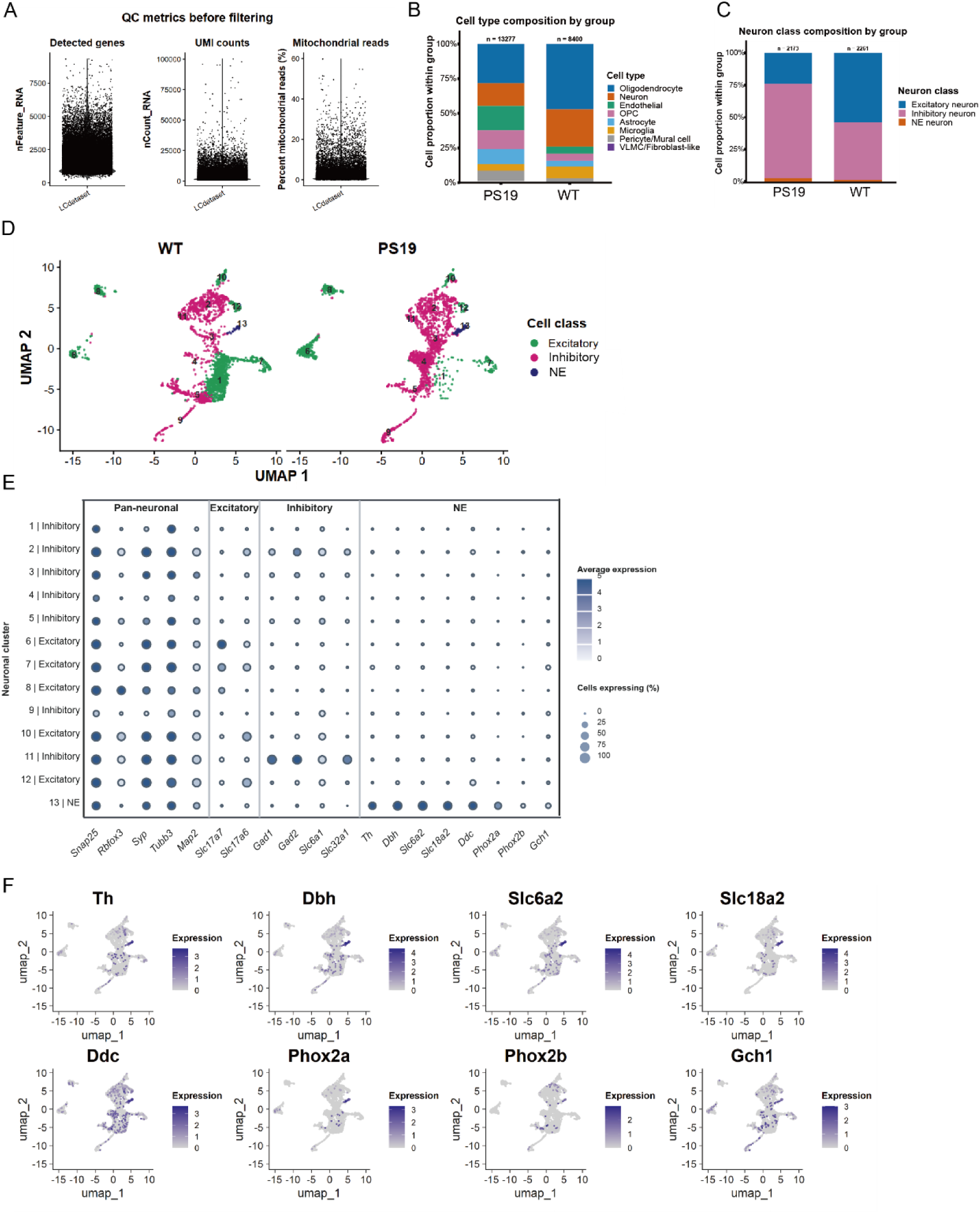
Quality control, cell composition, and marker expression in the LC single-cell data. (A) Quality-control metrics before filtering, showing the number of detected genes, UMI counts, and percentage of mitochondrial reads per barcode. (B) Proportions of major cell types in PS19 and WT samples; numbers above bars indicate total cells (PS19, n = 13277 cells; WT, n = 8400 cells). (C) Composition of neuronal classes within each group (PS19, n = 2173; WT, n = 2261). (D) UMAP of the neuronal subset, showing clusters and their excitatory, inhibitory, or noradrenergic annotations. (E) Dot plot of canonical marker expression across neuronal clusters; dot size indicates the percentage of cells expressing each marker, and color indicates average expression. (F) UMAP feature plots showing expression of the indicated noradrenergic markers. (WT: n = 8 mice; PS19: n = 8 mice, eight mice per genotype were pooled into one sample per genotype).

**FIGURE S6.**
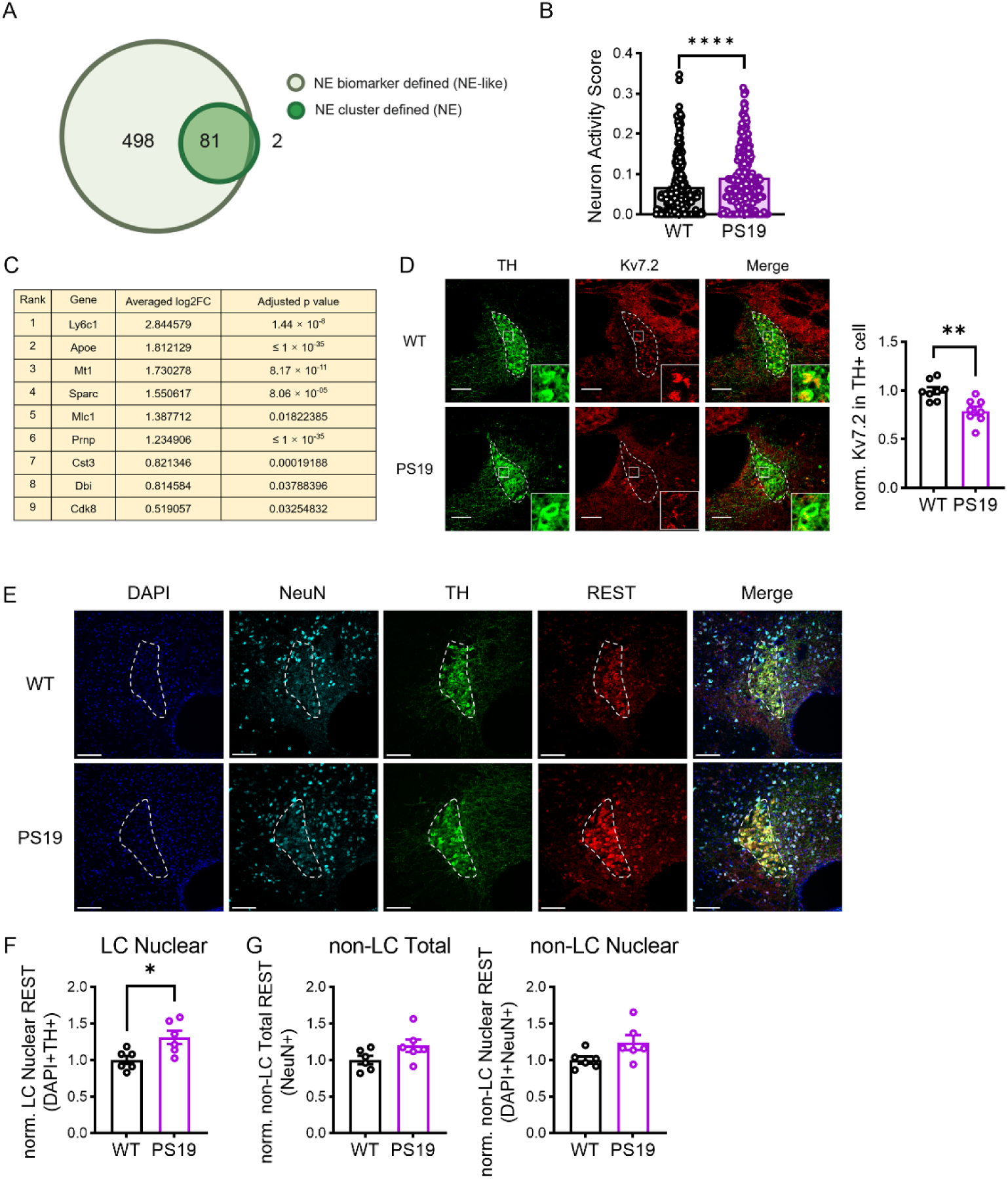
Comparison of NE neuron definitions and associated activity and molecular features. (A) Overlap between NE-like neurons, defined by detection of at least one of *Th, Dbh, Slc6a2,* or *Slc18a2*, and NE neurons in cluster 13. Numbers indicate cells unique to each group or shared by both. (B) Neuron activity scores in the NE-like population, compared between WT and PS19. (WT, n = 307 cells; PS19, n = 272 cells) (p < 0.0001) (C) The nine upregulated genes identified in the NE-like population, ranked by average log₂ fold change (PS19/WT), with adjusted P values. (D) Representative LC images showing TH (green), Kv7.2 (red), and merged staining in WT and PS19 mice, with quantification. (WT, n = 8 mice; PS19, n = 8 mice). (p = 0.0047) (E-G) Representative LC images showing DAPI (blue), NeuN (cyan), TH (green), REST (red), merged staining (E) and quantification (F-G). Dashed outlines indicate the LC; (Scale bars,100 μ m) (WT, n = 6 mice; PS19, n = 6 mice). (F: p = 0.0260) Data are presented as [mean ± SEM] in Table S5. Mann–Whitney tests were performed.

**FIGURE S7.**
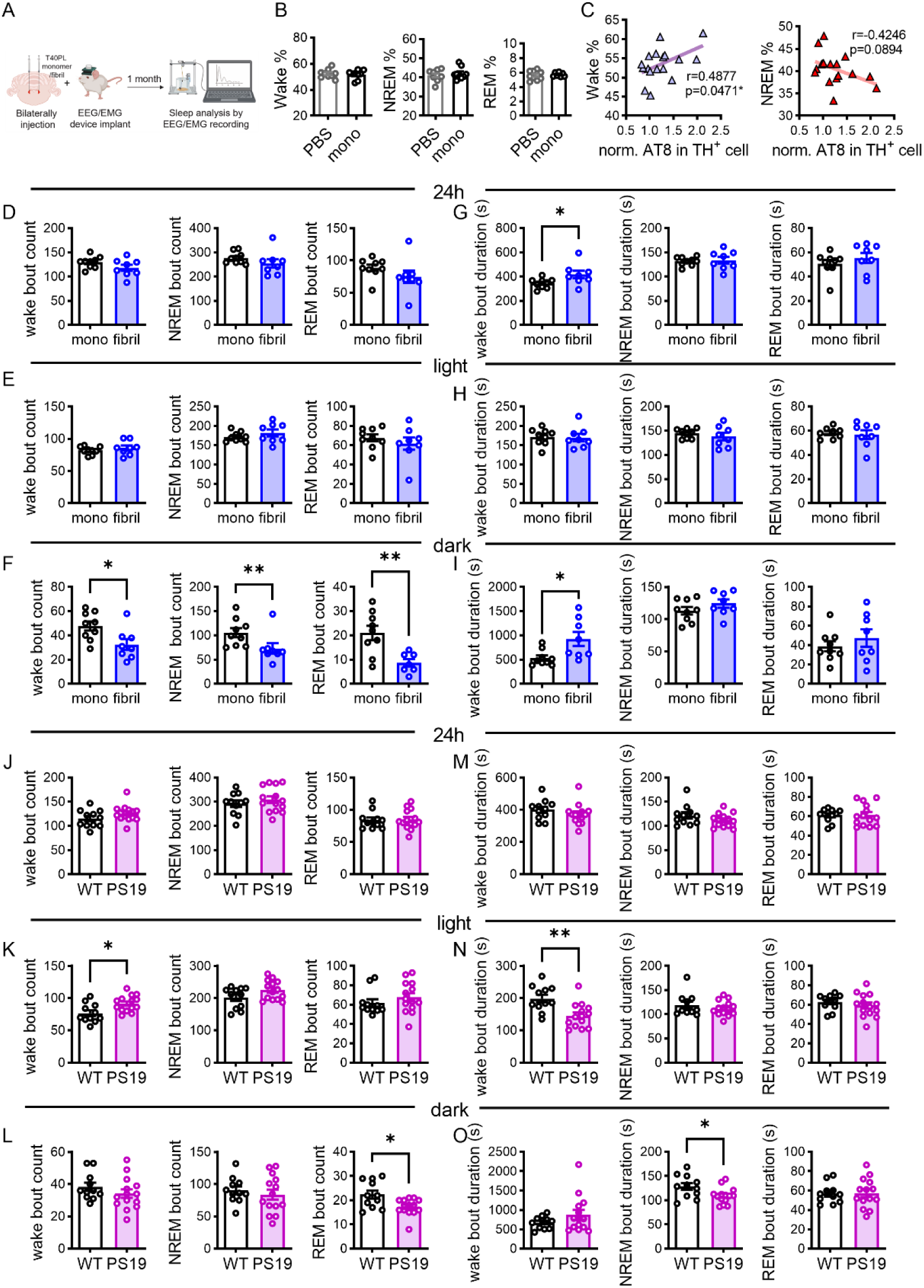
Sleep structure of tau fibril-inoculated mice and young PS19 mice. (A) Schematic illustration of the experimental design of LC inoculation of tau monomer or fibril followed by EEG/EMG assessment at 1-month timepoint. (B) Tau monomer did not change the percentages of time spent in wake (left), NREM (middle), and REM (right) during the 24-hour day compared to the PBS-inoculated wild-type mice. (PBS: n = 9 mice; mono: n = 9 mice) (C) AT8 signal of TH-positive cells in LC positively correlated with wake durations (left) and negatively correlated with NREM durations (right) during the 24-hour day. (n = 17 mice, containing both monomer and fibril inoculated wild-type mice) (D–I) Sleep structure of bout count (D-F) and mean bout duration (G-I) in 24-hour (upper panels), light cycle (middle panels) and dark cycle (below panels) of monomer- and fibril- inoculated mice. Left panels, wake, Middle panels, NREM, Right panels, REM. (mono: n = 9 mice; fibril: n = 8 mice) (J-O) Sleep structure of bout count (J-L) and mean bout duration (M-O) in 24-hour (upper panels), light cycle (middle panels) and dark cycle (below panels) of 2-month-old PS19 and WT mice. Left panels, wake, Middle panels, NREM, Right panels, REM. (WT: n = 11 mice; PS19: n = 14 mice) Data shown in Table S7. Mann-Whitney U test was used for unpaired data. Pearson correlation coefficient was calculated for correlation analysis.

**FIGURE S8.**
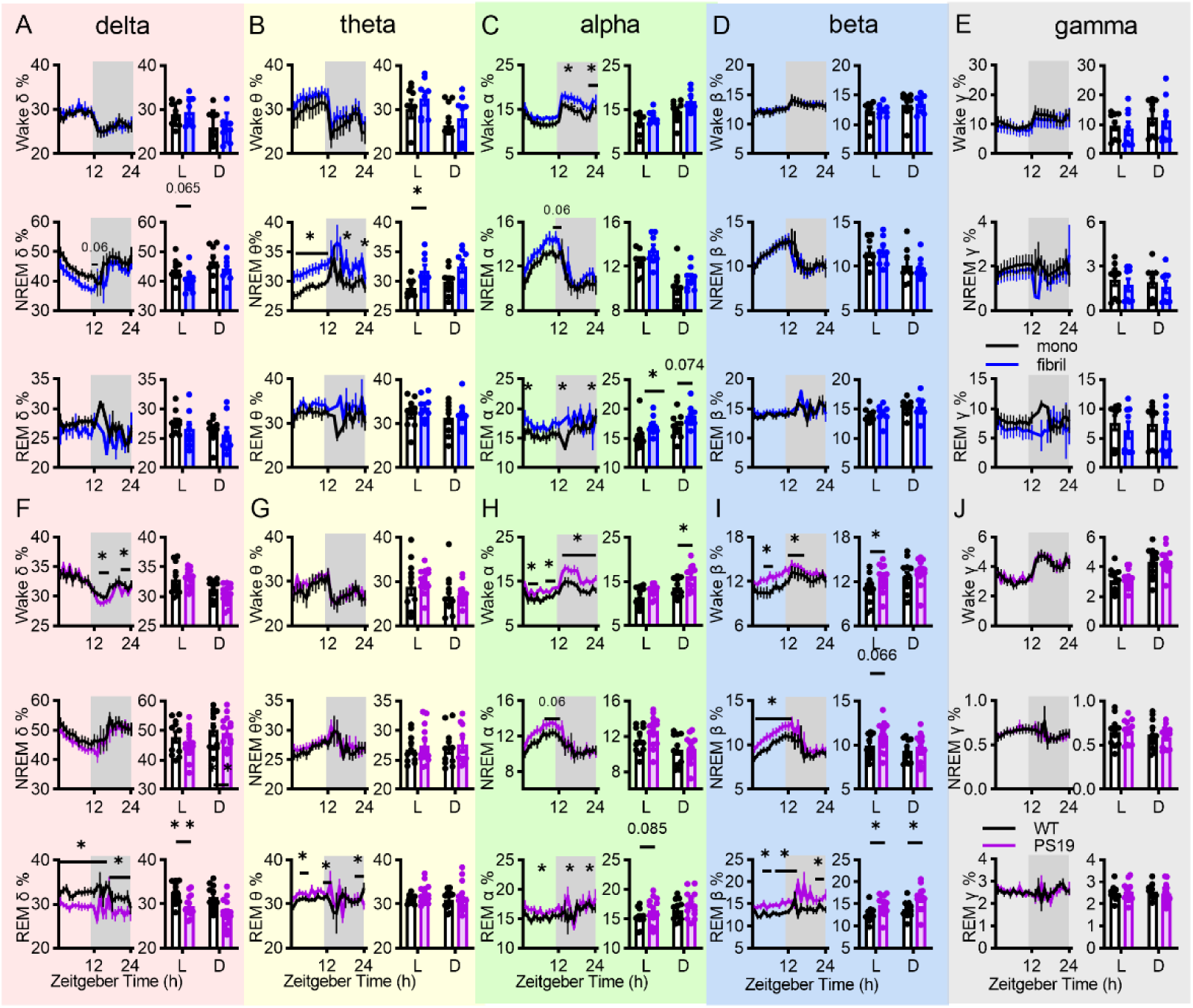
State-resolved temporal profiles of EEG-band power. (A–E) Comparisons between monomer and fibril-inoculated wild-type mice in the δ (A), θ (B), α (C), β (D), and γ (E) frequency bands (mono: n = 9 mice; fibril: n = 8 mice). Each panel contains hourly band-power profiles during wakefulness (top panels), NREM sleep (middle panels), and REM sleep (bottom panels), with mean values during the light (L) and dark (D) phases. (F–J) Comparisons between WT and PS19 mice in the δ (F), θ (G), α (H), β (I), and γ (J) frequency bands (WT: n = 11 mice; PS19: n = 14 mice). Each panel contains hourly band-power profiles during wakefulness (top panels), NREM sleep (middle panels), and REM sleep (bottom panels), with mean values during the light (L) and dark (D) phases. Multiple t-test were used to analysis power spectrum in light/dark phase.

**FIGURE S9.**
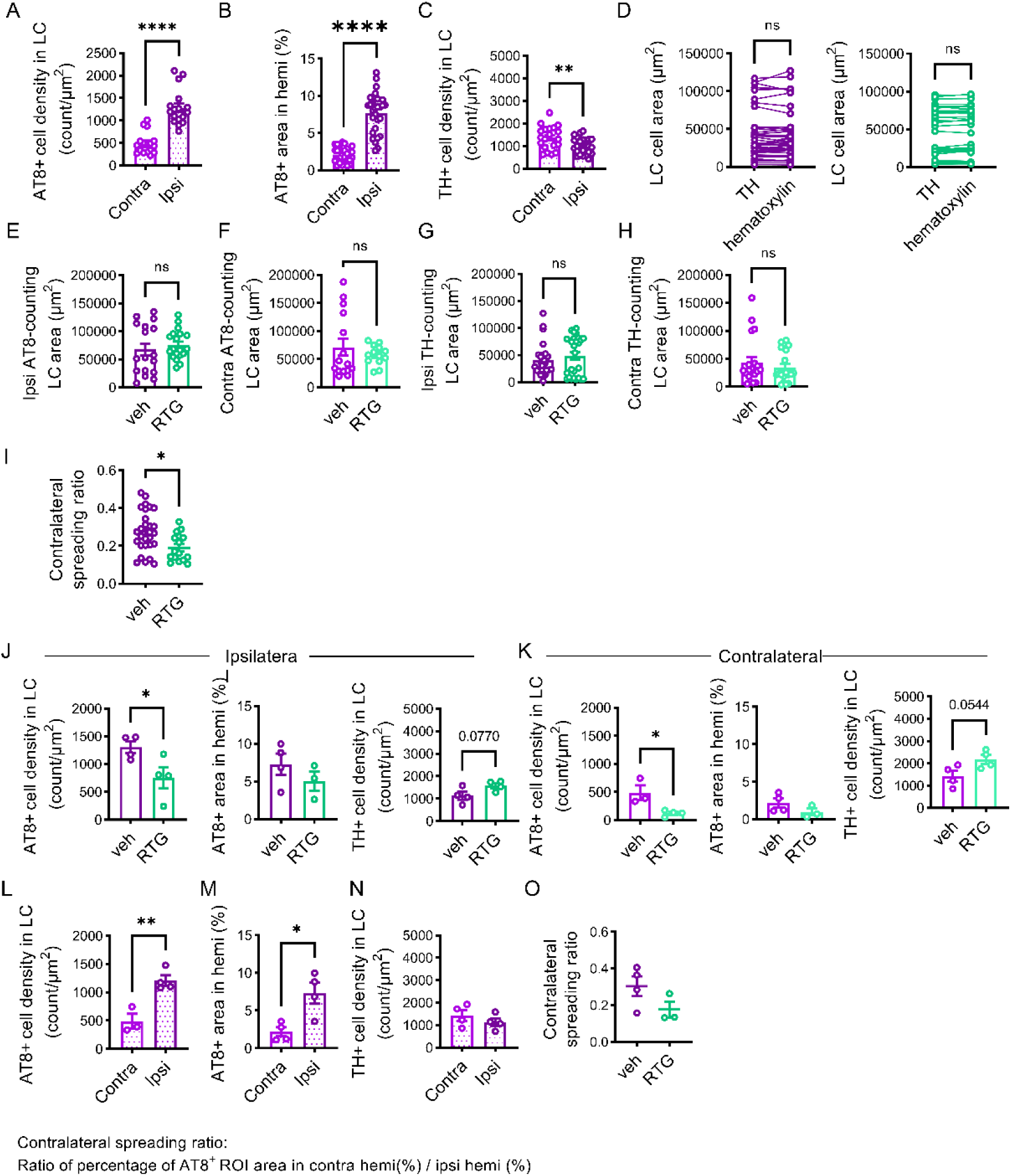
Section- and mouse-level analyses of AT8 burden and TH-positive cell density following unilateral LC fibril inoculation and retigabine treatment in PS19 mice. (A–I) Section-level analyses. (A–C) Ipsilateral versus contralateral LC AT8-positive cell density (A), hemispheric AT8-positive area fraction (B), and LC TH-positive cell density (C) in vehicle- treated mice (A, p < 0.0001; B, p < 0.0001; C, p = 0.0068). (D) Paired comparison of LC areas delineated using hematoxylin or TH staining. (E–H) LC ROI areas in vehicle- and retigabine-treated mice for ipsilateral (E, G) and contralateral (F, H) AT8 (E, F) and TH (G, H) analyses. (I) Contralateral-to-ipsilateral ratio of hemispheric AT8-positive area fractions in vehicle- and retigabine-treated mice (p = 0.0079). (J–O) Mouse-level analyses using measurements averaged across sections from each animal (unpaired t-test). (J, K) Ipsilateral (p = 0.0462) (J) and contralateral (p = 0.0225) (K) LC AT8- positive cell density (left), hemispheric AT8-positive area fraction (middle), and LC TH-positive cell density (right) in vehicle- and retigabine-treated mice. (L–N) Ipsilateral versus contralateral LC AT8-positive cell density (p = 0.0054) (L), hemispheric AT8-positive area fraction (p = 0.0154) (M), and LC TH-positive cell density (N) in vehicle-treated mice. (O) Contralateral-to-ipsilateral ratio of hemispheric AT8-positive area fractions. Each point represents one section in A–I or one mouse in J–O. Data are presented as mean ± SEM; numerical summaries are provided in Table S8. Paired comparisons used two-sided Wilcoxon signed-rank tests; unpaired comparisons used two-sided Mann–Whitney U tests. Otherwise mentioned in the legends.

**Supplementary Table 1.** Key resources table.

| **Antibodies** | SOURCE | IDENTIFIER |
|---|---|---|
| AT8 | Thermo Fisher | MN1020 |
| tau | Dako | A0024 |
| actin | Beyotime | AF0003 |
| tubulin | MBL | PM054 |
| TH | Millipore | AB152 |
| TH | Millipore | AB1542 |
| NeuN | Millipore | ABN91 |
| GFAP | Dako | Z0334 |
| Iba1 | wako | 019-19741 |
| Kv7.2 (KCNQ2) | Abcam | Ab22897 |
| Kv7.3 (KCNQ3) | Alomone Labs | APC-051 |
| REST | Proteintech | 22242-1-AP |
| **Reagents** |  |  |
| Tetrodotoxin (TTX) | Sigma-Aldrich | Cat# 4368-28-9 |
| DL-APV | Tocris | Cat# 0105 |
| SR-95531 (gabazine) | Tocris | Cat# 1262 |
| CNQX | Tocris | Cat# 0190 |
| Adensine 5’-triphosphate magnesium salt (MgATP) | Sigma-Aldrich | Cat# A9187 |
| Guanosine 5’-triphosphate sodium salt (Na<sub>3</sub>GTP) | Sigma-Aldrich | Cat# G8877 |
| Phosphocreatine disodium salt hydrate (Na-phosphocreatine) | Sigma-Aldrich | Cat# P7936 |
| Retigabine | Selleck | Cat# S4733 |
| AR-A014418 | MedChemExpress | HY-10512 |
| NS5806 | MedChemExpress | HY108588 |
| **Experimental Models: Strains** |  |  |
| PS19 mouse: | The Jackson Laboratory | RRID: 034848 |
| Wild type mouse: *C57BL/6J* | The Jackson Laboratory | RRID: 000664 |
| mTauKO: *B6.129X1-Mapt<sup>tm1Hnd</sup>/J* | The Jackson Laboratory | RRID:007251 |
| **Software** |  |  |
| NeuroScore | Data Sciences Internationial | RRID:SCR_017107 |
| Prism | GraphPad | RRID:002798 |
| Mini Analysis | Synaptosoft | RRID:002184 |
| MATLAB R2016a | MathWorks | RRID:001622 |
| BioRender | BioRender | RRID:SCR_018361 |
| Fiji ImageJ | SciCrunch Registry | RRID:SCR_003070 |
| Imaris | Oxford Instruments | RRID:SCR_007370 |
| QuPath | University of Edinburg | RRID:SCR_018257 |

**Supplementary Table 2.** Data in Fig 1 and Fig S1.

| Fig Location | 10M WT (ave ± SEM) | 10M PS19 (ave ± SEM) | P value |
|---|---|---|---|
| Fig 1C | 1192.82 ± 108.05 count/mm<sup>2</sup> | 782.93 ± 39.52 count/mm<sup>2</sup> | 0.0286 |
| Fig 1D | 1.00 ± 0.18 | 5.22 ± 0.35 | 0.0286 |
| Fig 1G left | 6.02 ± 1.23 Hz | 5.78 ± 0.91 Hz | >0.9999 |
| Fig 1G right | -46.67 ± 0.86 mV | -47.26 ± 0.98 mV | 0.6989 |
| Fig S1A | 1.00 ± 0.09 | 1.65 ± 0.19 | 0.1000 |
| Fig S1B | 142.24 ± 15.06 count/mm<sup>2</sup> | 202.15 ± 34.59 count/mm<sup>2</sup> | 0.4000 |
| Fig Location | 2M WT (ave ± SEM) | 2M PS19 (ave ± SEM) | P value |
| Fig 1K | 1373.85 ± 31.84 count/mm<sup>2</sup> | 1292.09 ± 70.12 count/mm<sup>2</sup> | 0.5476 |
| Fig 1L | 1.00 ± 0.04 | 5.31 ± 0.47 | 0.0079 |
| Fig 1O left | 5.66 ± 0.54 Hz | 8.76 ± 0.62 Hz | 0.0012 |
| Fig 1O right | -46.46 ± 0.76 mV | -48.26 ± 0.83 mV; | 0.1382 |
| Fig S1C | 1.00 ± 0.09 | 0.99 ± 0.14 | 0.6200 |
| Fig S1D | 147 ± 15.26 count/mm<sup>2</sup> | 141.37 ±12.02 count/mm<sup>2</sup> | 0.9015 |

**Supplementary Table 3.**
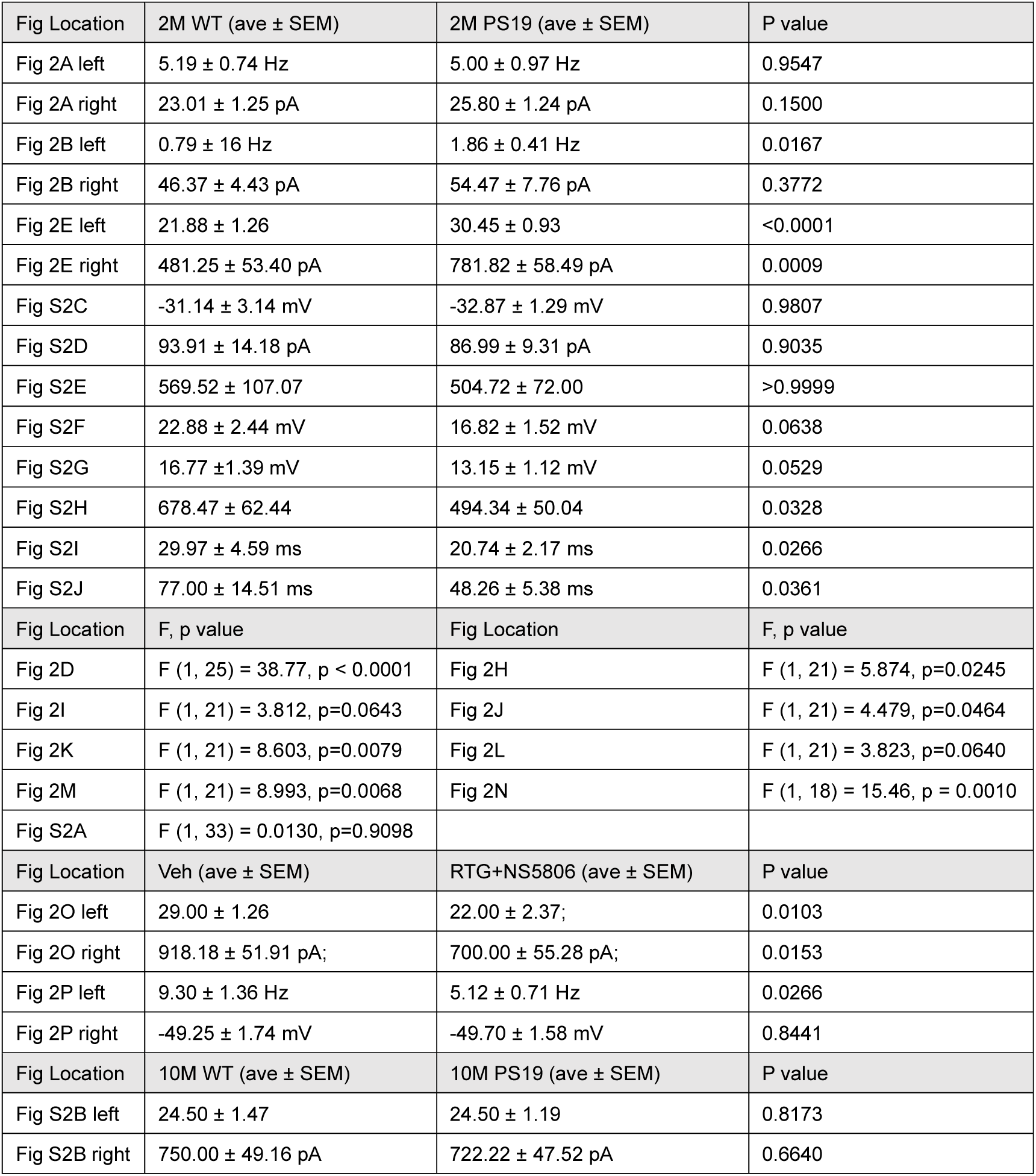
Data in Fig 2 and Fig S2.

**Supplementary Table 4.** Data in Fig 3 and Fig S3 and Fig S4.

| Fig Location | mono (ave ± SEM) | fibril (ave ± SEM) | P value |
|---|---|---|---|
| Fig 3C left | 1.00 ± 0.06 | 1.42 ± 0.11 | 0.0047 |
| Fig 3C right | 53.77 ± 0.91 | 69.87 ± 3.86 | 0.0027 |
| Fig 3D | 1.00 ± 0.27 | 1.99 ± 0.21 | 0.0317 |
| Fig 3G | 1.00 ± 0.05 | 1.52 ± 0.13 | 0.0006 |
| Fig 3J left | 6.38 ± 0.73 Hz | 10.11 ± 0.80 Hz | 0.0049 |
| Fig 3J right | -46.89 ± 1.44 mV | -47.92 ± 1.00 mV | 0.4954 |
| Fig 3M left | 22.85 ± 1.65 | 28.16 ± 1.10 | 0.0470 |
| Fig 3M right | 553.85 ± 53.85 pA | 794.74 ± 58.48 pA | 0.0117 |
| Fig S3B left | 961.19 ± 46.80 | 893.06 ± 37.02 | 0.4649 |
| Fig S3B right | 1.00 ± 0.05 | 0.98 ± 0.06 | >0.9999 |
| Fig S3C | 1.00 ± 0.15 | 1.13 ± 0.19 | 0.7922 |
| Fig S3D | 1.00 ± 0.02 | 0.99 ± 0.06 | 0.7922 |
| Fig S3E | 1.00 ± 0.07 | 1.21 ± 0.05 | 0.0823 |
| Fig S3F | 1.00 ± 0.05 | 1.21 ± 0.07 | 0.0823 |
| Fig S3G | 1.00 ± 0.04 | 1.00 ± 0.09 | 0.7922 |
| Fig S3H | 1.00 ± 0.06 | 1.06 ± 0.09 | 0.7922 |
| Fig S3I left | 30.57 ± 4.48 | 32.67 ± 2.79 | 0.3497 |
| Fig S3I right | 614.29 ± 45.92 pA | 700.00 ± 51.64 pA | 0.2552 |
| Fig S4A | -35.84 ± 1.02 mV | -36.58 ± 0.80 mV | 0.7550 |
| Fig S4B | 72.65 ± 9.87 pA | 69.84 ± 4.95 pA | 0.8723 |
| Fig S4C | 753.09 ± 95.68 | 598.63 ± 70.03 | 0.1549 |
| Fig Location | F, p value | Fig Location | F, p value |
| Fig 3O left | F (1, 23) = 2.540, p=0.1246 | Fig 3O right | F (1, 23) = 4.836, p=0.0382 |
| Fig 3P left | F (1, 23) = 2.994, p=0.0970 | Fig 3P right | F (1, 23) = 6.850, p=0.0154 |
| Fig S4D left | F (1, 23) = 1.382, p=0.2517 | Fig S4D right | F (1, 23) = 1.637, p=0.2135 |
| Fig 3Q | F (1, 16) = 23.75, p=0.0002 | Fig S3I | F (1, 11) = 0.0399, p=0.8453 |
| Fig S4F | F (1, 22) = 1.304, p=0.2658 | Fig 3L | F (1, 30) = 17.87, p = 0.0002 |
| Fig Location | Veh (ave ± SEM) | RTG+NS5806 (ave ± SEM) | P value |
| Fig 3R left | 27.13 ± 1.27 | 18.30 ± 1.37 | 0.0004 |
| Fig 3R right | 787.50 ± 39.81 pA | 510.00 ± 70.63 pA | 0.0104 |
| Fig 3S left | 9.47 ± 0.82 Hz | 3.86 ± 0.39 Hz | <0.0001 |
| Fig 3S right | -49.02 ± 1.35 mV | -51.21 ± 0.79 mV | 0.0727 |
| Fig Location | PBS (ave ± SEM) | mono (ave ± SEM) | P value |
| Fig S3A | 1.00 ± 0.02 | 0.93 ± 0.05 | 0.2973 |
| Fig Location | mTauKO + mono (ave ± SEM) | mTauKO + fibril (ave ± SEM) | P value |
| Fig S4F left | 32.83 ± 2.16 | 33.75 ± 1.23 | 0.6380 |
| Fig S4F right | 683.33 ± 47.41 pA | 616.67 ± 38.60 pA | 0.4793 |
| Fig S4G left | -44.36 ± 1.78 mV | -41.55 ± 1.33 mV | 0.1925 |
| Fig S4G middle | 66.13 ± 7.96 pA | 51.41 ± 6.65 pA | 0.1925 |
| Fig S4G right | 899.03 ± 234.44 | 1119.03 ± 182.40 | 0.0616 |

**Supplementary Table 5.** Data in Fig 4, Fig 5 and Fig S6.

| Fig Location | Veh (ave ± SEM) | AR (ave ± SEM) | P value |
|---|---|---|---|
| Fig 4F left | 28.71 ± 0.62 | 21.44 ± 2.21 | 0.0027 |
| Fig 4F right | 721.43 ± 48.24 pA | 387.50 ± 39.82 pA | < 0.0001 |
| Fig 4I left | 9.99 ± 0.94 Hz | 6.25 ± 1.07 Hz | 0.0137 |
| Fig 4I right | -48.29 ± 0.83 mV | -46.60 ± 1.26 mV | 0.1524 |
| Fig Location | F, p value | Fig Location | F, p value |
| Fig 4B left | F (1, 24) = 2.039, p=0.1662 | Fig 4B middle | F (1, 24) = 0.9391, p=0.3422 |
| Fig 4B right | F (1, 24) = 4.982 p=0.0352 | Fig 4C left | F (1, 24) = 1.496, p=0.2332 |
| Fig 4C middle | F (1, 24) = 0.6036, p=0.4448 | Fig 4C right | F (1, 24) = 5.364, p=0.0294 |
| Fig 4E | F (1, 28) = 27.13, p < 0.0001 |  |  |
| Fig Location | 2M WT (ave ± SEM) | 2M PS19 (ave ± SEM) | P value |
| Fig 5D | 0.03 ± 0.01 | 0.07 ± 0.01 | 0.0008 |
| Fig 5H | 1.00 ± 0.06 | 1.29 ± 0.09 | 0.0152 |
| Fig 5I | 1.00 ± 0.02 | 0.72 ± 0.05 | 0.0070 |
| Fig S6B | 0.068 ± 0.004 | 0.091 ± 0.005 | <0.0001 |
| Fig S6D | 1.00 ± 0.04 | 0.79 ± 0.04 | 0.0047 |
| Fig S6F | 1.00 ± 0.06 | 1.31 ± 0.09 | 0.0260 |
| Fig S6G left | 1.00 ± 0.06 | 1.20 ± 0.09 | 0.1320 |
| Fig S6G right | 1.00 ± 0.05 | 1.24 ± 0.10 | 0.0931 |

**Supplementary Table 6.**
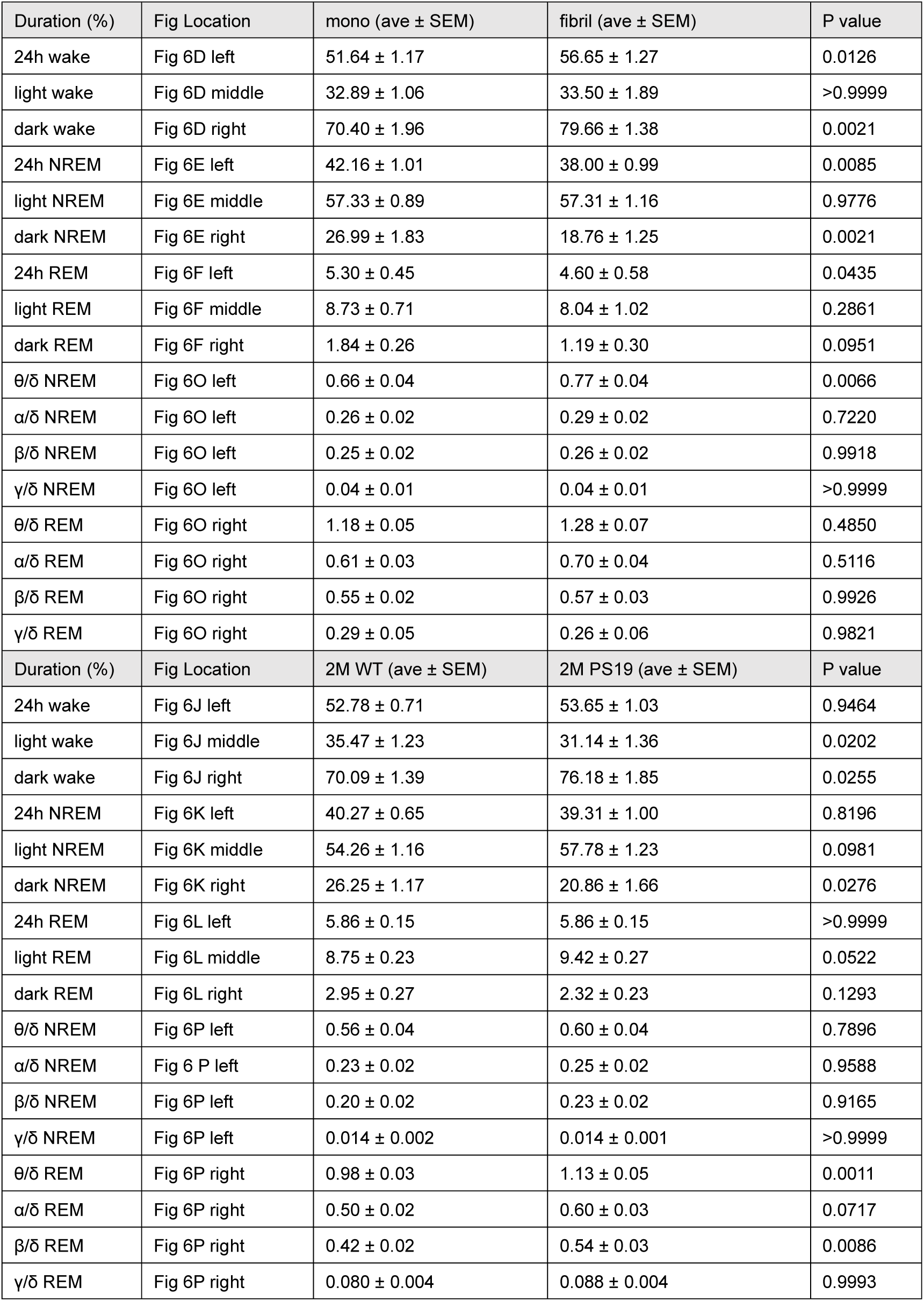
Data in Fig 6.

| Duration (%) | Fig Location | mono (ave ± SEM) | fibril (ave ± SEM) | P value |
|---|---|---|---|---|
| 24h wake | Fig 6D left | 51.64 ± 1.17 | 56.65 ± 1.27 | 0.0126 |
| light wake | Fig 6D middle | 32.89 ± 1.06 | 33.50 ± 1.89 | >0.9999 |
| dark wake | Fig 6D right | 70.40 ± 1.96 | 79.66 ± 1.38 | 0.0021 |
| 24h NREM | Fig 6E left | 42.16 ± 1.01 | 38.00 ± 0.99 | 0.0085 |
| light NREM | Fig 6E middle | 57.33 ± 0.89 | 57.31 ± 1.16 | 0.9776 |
| dark NREM | Fig 6E right | 26.99 ± 1.83 | 18.76 ± 1.25 | 0.0021 |
| 24h REM | Fig 6F left | 5.30 ± 0.45 | 4.60 ± 0.58 | 0.0435 |
| light REM | Fig 6F middle | 8.73 ± 0.71 | 8.04 ± 1.02 | 0.2861 |
| dark REM | Fig 6F right | 1.84 ± 0.26 | 1.19 ± 0.30 | 0.0951 |
| θ/δ NREM | Fig 6O left | 0.66 ± 0.04 | 0.77 ± 0.04 | 0.0066 |
| α/δ NREM | Fig 6O left | 0.26 ± 0.02 | 0.29 ± 0.02 | 0.7220 |
| β/δ NREM | Fig 6O left | 0.25 ± 0.02 | 0.26 ± 0.02 | 0.9918 |
| γ/δ NREM | Fig 6O left | 0.04 ± 0.01 | 0.04 ± 0.01 | >0.9999 |
| θ/δ REM | Fig 6O right | 1.18 ± 0.05 | 1.28 ± 0.07 | 0.4850 |
| α/δ REM | Fig 6O right | 0.61 ± 0.03 | 0.70 ± 0.04 | 0.5116 |
| β/δ REM | Fig 6O right | 0.55 ± 0.02 | 0.57 ± 0.03 | 0.9926 |
| γ/δ REM | Fig 6O right | 0.29 ± 0.05 | 0.26 ± 0.06 | 0.9821 |
| Duration (%) | Fig Location | 2M WT (ave ± SEM) | 2M PS19 (ave ± SEM) | P value |
| 24h wake | Fig 6J left | 52.78 ± 0.71 | 53.65 ± 1.03 | 0.9464 |
| light wake | Fig 6J middle | 35.47 ± 1.23 | 31.14 ± 1.36 | 0.0202 |
| dark wake | Fig 6J right | 70.09 ± 1.39 | 76.18 ± 1.85 | 0.0255 |
| 24h NREM | Fig 6K left | 40.27 ± 0.65 | 39.31 ± 1.00 | 0.8196 |
| light NREM | Fig 6K middle | 54.26 ± 1.16 | 57.78 ± 1.23 | 0.0981 |
| dark NREM | Fig 6K right | 26.25 ± 1.17 | 20.86 ± 1.66 | 0.0276 |
| 24h REM | Fig 6L left | 5.86 ± 0.15 | 5.86 ± 0.15 | >0.9999 |
| light REM | Fig 6L middle | 8.75 ± 0.23 | 9.42 ± 0.27 | 0.0522 |
| dark REM | Fig 6L right | 2.95 ± 0.27 | 2.32 ± 0.23 | 0.1293 |
| θ/δ NREM | Fig 6P left | 0.56 ± 0.04 | 0.60 ± 0.04 | 0.7896 |
| α/δ NREM | Fig 6 P left | 0.23 ± 0.02 | 0.25 ± 0.02 | 0.9588 |
| β/δ NREM | Fig 6P left | 0.20 ± 0.02 | 0.23 ± 0.02 | 0.9165 |
| γ/δ NREM | Fig 6P left | 0.014 ± 0.002 | 0.014 ± 0.001 | >0.9999 |
| θ/δ REM | Fig 6P right | 0.98 ± 0.03 | 1.13 ± 0.05 | 0.0011 |
| α/δ REM | Fig 6P right | 0.50 ± 0.02 | 0.60 ± 0.03 | 0.0717 |
| β/δ REM | Fig 6P right | 0.42 ± 0.02 | 0.54 ± 0.03 | 0.0086 |
| γ/δ REM | Fig 6P right | 0.080 ± 0.004 | 0.088 ± 0.004 | 0.9993 |

**Supplementary Table 7.** Data in Fig S7.

| Fig Location | PBS (ave ± SEM) | mono (ave ± SEM) | P value |
|---|---|---|---|
| Fig S7B left | 52.30 ± 1.14 | 51.64 ± 1.17 | 0.8814 |
| Fig S7B middle | 41.10 ± 1.02 | 42.16 ± 1.01 | 0.7149 |
| Fig S7B right | 5.61 ± 0.21 | 5.74 ± 0.09 | 0.7598 |
| Fig Location | mono (ave ± SEM) | fibril (ave ± SEM) | P value |
| Fig S7D left | 129.89 ± 4.02 | 118.38 ± 6.39 | 0.1391 |
| Fig S7D middle | 277.22 ± 8.09 | 255.13 ± 17.66 | 0.2553 |
| Fig S7D right | 88.67 ± 5.04 | 74.88 ± 9.62 | 0.2089 |
| Fig S7E left | 82.11 ± 2.05 | 85.88 ± 3.90 | 0.6520 |
| Fig S7E middle | 171.78 ± 4.17 | 182.00 ± 8.91 | 0.4100 |
| Fig S7E right | 67.67 ± 3.50 | 61.75 ± 6.37 | 0.4084 |
| Fig S7F left | 47.78 ± 3.87 | 32.50 ± 4.46 | 0.0192 |
| Fig S7F middle | 105.44 ± 9.46 | 73.13 ± 10.74 | 0.0054 |
| Fig S7F right | 21.00 ± 2.98 | 8.71 ± 1.52 | 0.0072 |
| Fig S7G left | 345.21 ± 13.47 | 417.15 ± 30.79 | 0.0360 |
| Fig S7G middle | 131.83 ± 3.00 | 133.89 ± 6.81 | 0.7620 |
| Fig S7G right | 50.71 ± 3.14 | 55.40 ± 4.16 | 0.3704 |
| Fig S7H left | 172.14 ± 7.06 | 169.84 ± 9.55 | 0.6730 |
| Fig S7H middle | 144.70 ± 3.28 | 138.04 ± 7.65 | 0.4807 |
| Fig S7H right | 58.13 ± 1.60 | 56.83 ± 3.52 | 0.9591 |
| Fig S7I left | 530.99 ± 54.38 | 926.59 ± 145.10 | 0.0111 |
| Fig S7I middle | 113.28 ± 5.80 | 124.44 ± 6.47 | 0.1627 |
| Fig S7I right | 38.80 ± 5.26 | 47.36 ± 8.96 | 0.6058 |
| Fig Location | 2M WT (ave ± SEM) | 2M PS19 (ave ± SEM) | P value |
| Fig S7J left | 115.18 ± 5.39 | 125.79 ± 4.56 | 0.2274 |
| Fig S7J middle | 292.09 ± 13.66 | 308.57 ± 13.83 | 0.5357 |
| Fig S7J right | 84.09 ± 4.11 | 84.93 ± 4.04 | 0.9037 |
| Fig S7K left | 76.82 ± 4.39 | 91.64 ± 3.31 | 0.0122 |
| Fig S7K middle | 201.55 ± 9.89 | 224.79 ± 7.99 | 0.1492 |
| Fig S7K right | 61.73 ± 3.84 | 67.64 ± 4.26 | 0.2167 |
| Fig S7L left | 38.36 ± 2.53 | 34.14 ± 2.65 | 0.2166 |
| Fig S7L middle | 90.55 ± 6.05 | 83.79 ± 7.80 | 0.4254 |
| Fig S7L right | 22.36 ± 1.67 | 17.29 ± 0.92 | 0.0308 |
| Fig S7M left | 400.17 ± 18.90 | 372.61 ± 17.17 | 0.2225 |
| Fig S7M middle | 121.75 ± 6.41 | 111.84 ± 3.77 | 0.2915 |
| Fig S7M right | 61.04 ± 2.10 | 61.26 ± 3.00 | 0.7267 |
| Fig S7N left | 198.73 ± 12.08 | 145.86 ± 10.12 | 0.0028 |
| Fig S7N middle | 119.10 ± 6.79 | 112.56 ± 4.09 | 0.6867 |
| Fig S7N right | 62.63 ± 2.44 | 59.09 ± 3.17 | 0.4427 |
| Fig S7O left | 674.08 ± 48.24 | 878.36 ± 126.83 | 0.3730 |
| Fig S7O middle | 128.35 ± 7.05 | 109.40 ± 4.44 | 0.0442 |
| Fig S7O right | 56.91 ± 3.09 | 57.47 ± 4.05 | 0.8508 |

**Supplementary Table 8.** Data in Fig 7 and Fig S9.

| Fig Location | veh (ave ± SEM) | RTG (ave ± SEM) | P value |
|---|---|---|---|
| Fig 7F left | 1287.02 ± 95.20 count/µm<sup>2</sup> | 749.27 ± 89.55 count/µm<sup>2</sup> | 0.0002 |
| Fig 7F middle | 7.61 ± 0.58 % | 6.33 ± 0.35 % | 0.0077 |
| Fig 7F right | 1006.07 ± 71.01 count/µm<sup>2</sup> | 1589.92 ± 87.05 count/µm<sup>2</sup> | <0.0001 |
| Fig 7G left | 486.08 ± 65.37 count/µm<sup>2</sup> | 122.06 ± 18.27 count/µm<sup>2</sup> | <0.0001 |
| Fig 7G middle | 2.10 ± 0.20 % | 1.15 ± 0.18 % | 0.0051 |
| Fig 7G right | 1399.27 ± 108.94 count/µm<sup>2</sup> | 2329.13 ± 171.87 count/µm<sup>2</sup> | <0.0001 |
| Fig S9E | 67543.52 ± 10753.91 µm<sup>2</sup> | 75830.55 ± 5944.50 µm<sup>2</sup> | 0.3969 |
| Fig S9F | 71403.59 ± 14943.60 µm<sup>2</sup> | 57804.42 ± 4857.85 µm<sup>2</sup> | 0.6177 |
| Fig S9G | 40833.73 ± 6070.29 µm<sup>2</sup> | 48620.05 ± 7096.68 µm<sup>2</sup> | 0.7437 |
| Fig S9H | 43374.95 ± 9465.15 µm<sup>2</sup> | 34673.39 ± 6527.00 µm<sup>2</sup> | 0.3314 |
| Fig S9I | 0.28 ± 0.02 | 0.19 ± 0.02 | 0.0174 |
| Fig Location | Contra (ave ± SEM) | Ipsi (ave ± SEM) | P value |
| Fig S9A | 486.08 ± 65.37 count/µm<sup>2</sup> | 1287.02 ± 95.20 count/µm<sup>2</sup> | <0.0001 |
| Fig S9B | 2.10 ± 0.20 % | 7.67 ± 0.51 % | <0.0001 |
| Fig S9C | 1374.74 ± 114.71 count/µm<sup>2</sup> | 1006.07 ± 71.01 count/µm<sup>2</sup> | 0.0086 |
| Fig Location | TH (ave ± SEM) | hematoxylin (ave ± SEM) | P value |
| Fig S9D left | 39983.87 ± 4647.09 µm<sup>2</sup> | 40206.75 ± 4704.34 µm<sup>2</sup> | 0.7205 |
| Fig S9D right | 41686.59 ± 4830.75 µm<sup>2</sup> | 41998.36 ± 4789.30 µm<sup>2</sup> | 0.2680 |
| Fig Location | veh (ave ± SEM) | RTG (ave ± SEM) | P value |
| Fig S9J left | 1304.73 ± 104.54 count/µm<sup>2</sup> | 754.54 ± 193.20 count/µm<sup>2</sup> | 0.0462 |
| Fig S9J middle | 7.31 ± 1.41 % | 5.05 ± 1.31 % | 0.3087 |
| Fig S9J right | 1134.26 ± 176.14 count/µm<sup>2</sup> | 1579.66 ± 112.32 count/µm<sup>2</sup> | 0.0770 |
| Fig S9K left | 489.88 ± 128.87 count/µm<sup>2</sup> | 124.84 ± 25.62 count/µm<sup>2</sup> | 0.0225 |
| Fig S9K middle | 2.19 ± 0.60 % | 0.98 ± 0.46 % | 0.1951 |
| Fig S9K right | 1440.06 ± 240.61 count/µm<sup>2</sup> | 2185.05 ± 199.14 count/µm<sup>2</sup> | 0.0544 |
| Fig S9O | 0.30 ± 0.05 | 0.18 ± 0.04 | 0.1488 |
| Fig Location | Contra (ave ± SEM) | Ipsi (ave ± SEM) | P value |
| Fig S9L | 489.88 ± 128.87 count/µm<sup>2</sup> | 1208.33 ± 92.31 count/µm<sup>2</sup> | 0.0054 |
| Fig S9M | 2.19 ± 0.60 % | 7.31 ± 1.41 % | 0.0154 |
| Fig S9N | 1440.06 ± 240.61 count/µm<sup>2</sup> | 1134.26 ± 176.14 count/µm<sup>2</sup> | 0.3447 |

